# Proteolysis-targeting vaccines (PROTAVs) for robust combination immunotherapy of melanoma

**DOI:** 10.1101/2024.10.01.616069

**Authors:** Qiyan Wang, Ting Su, Furong Cheng, Shurong Zhou, Xiang Liu, Mi Wang, You Xu, Ri Tang, Shimiao Liao, Jordan Dailey, Guolan Xiao, Chunpeng Yang, Hanning Wen, Weijia Zheng, Bo Wen, Katarzyna M Tyc, Jinze Liu, Duxin Sun, Shaomeng Wang, Guizhi Zhu

## Abstract

Protein/peptide subunit vaccines are promising to promote the tumor therapeutic efficacy of immune checkpoint blockade (ICB). However, current protein/peptide vaccines elicit limited antitumor T cell responses, leading to suboptimal therapeutic efficacy. Here, we present proteolysis-targeting vaccines (PROTAVs) that facilitate antigen proteolytic processing and cross-presentation to potentiate T cell responses for robust ICB combination immunotherapy of melanoma. PROTAVs are modular conjugates of protein/peptide antigens, E3 ligase-binding ligands, and linkers. In antigen-presenting cells (APCs), PROTAVs bind to E3 ligases to rapidly ubiquitinate PROTAV antigens, facilitating antigen proteolytic processing by proteasome, and thereby promoting antigen cross-presentation to T cells and potentiating CD8^+^ T cell responses. We developed a melanoma PROTAV using a tandem peptide of trivalent melanoma-associated antigens. Co-delivered by lipid nanoparticles (LNPs) with bivalent immunostimulant adjuvants, this PROTAV promotes the quantity and quality of melanoma-specific CD8^+^ T cells in mice. Further, combining PROTAV and ICB ameliorates the immunosuppressive melanoma microenvironment. As a result, PROTAV and ICB combination enhances melanoma complete regression rates and eradicated 100% large *Braf^V600E^* melanoma without recurrence in syngeneic mice. PROTAVs hold the potential for robust tumor combination immunotherapy.

---

Cancer immunotherapy, such as ICB, has significantly improved the treatment outcomes for patients with many types of cancers, including melanoma.^1,2^ Yet, most advanced melanoma patients still do not durably respond to current immune checkpoint inhibitors (ICIs) or suffer from immune-related adverse effects (irAEs) due to imbalanced immune homeostasis.^3–5^ Cancer therapeutic vaccines hold great potential to elicit or expand antitumor immunity, making cancer vaccines appealing to improve the treatment outcome of ICB tumor immunotherapy.^6–10^ However, current cancer therapeutic vaccines have shown limited clinical efficacy,^7,11^ largely due to the overall poor immunogenicity of tumor-associated antigens (TAAs) and neoantigens.^12^ This demands innovative technologies to promote the ability of cancer vaccines to elicit antitumor immunity, especially antigen-specific CD8^+^ cytotoxic T lymphocytes (CTLs) that are central to the immunotherapy of cancer, including melanoma.

The ability of vaccines to elicit or expand CD8^+^ CTLs is hinged on the efficiency of antigen processing and cross-presentation by APCs.^13,14^ Specifically, in the cytosol of APCs [e.g., dendritic cells (DCs)], vaccine antigens undergo ubiquitination, allowing for antigen processing into antigenic minimal epitopes by 26S proteasome via proteolytic cleavage.^15^ The resulting minimal epitopes are then transported by transporter associated with antigen processing (TAP) from the cytosol into the endoplasmic reticulum, where minimal epitopes are complexed with major histocompatibility complex class I (MHC-I). Lastly, the MHC-epitope complexes are transported to APC cell surfaces, where epitopes are presented to CD8^+^ T cells to elicit CTL responses.^16^ The modulation of proteolytic cleavage kinetics of protein of interest has garnered tremendous attention over the past decade.^17–19^ Notably, proteolysis targeting chimeras (PROTACs) have been studied to facilitate the degradation of endogenous target proteins for various biomedical applications, such as cancer therapy.^1,2^ A PROTAC is comprised of a ligand for E3 ligase^21^ and a ligand for protein of interest (POI), which promotes the physical proximity of E3 ligase with POI, thereby facilitating the ubiquitination and proteolytic cleavage of the latter.^22^ Interestingly, recent evidence has emerged that PROTACs may promote antigen-specific immune responses against protein antigens^23^ and antigenic peptides derived from target proteins for PROTAC^24^. Therefore, we hypothesize that facilitating the proteolytic processing of exogenous vaccine antigens into minimal peptide epitopes in APCs may promote the corresponding antigen presentation and antigen-specific CD8^+^ CTL responses, thereby promoting their tumor therapeutic efficacy.

Here, we present PROTAVs as a vaccine platform that promotes antigen processing and cross-presentation in APCs, thereby promoting antitumor T cell responses for the combination immunotherapy of melanoma (**Fig. 1**). A PROTAV is a modular molecular conjugate of a peptide or protein antigen, an E3 ligase-binding ligand, and a linker between the antigen and the E3 ligand. The E3 ligand avidly binds to endogenous E3 ligase, thereby bringing E3 ligase into proximity of antigens to facilitate antigen ubiquitination and proteolytic processing by proteasome. In APCs, PROTAVs facilitated antigen proteolytic processing, and promoted antigen presentation, thus potentiating antigen-specific T cell response. Using a model antigen, we optimized the linker in PROTAV to elicit up to 3-fold antigen-specific CD8^+^ T cell response relative to unmodified vaccine. PROTAVs are broadly applicable to antigens of proteins and multivalent antigenic synthetic long peptides, both of which are common types of tumor antigens. For melanoma immunotherapy, we synthesized a trivalent PROTAV for a synthetic tandem peptide of three melanoma-associated antigens. In mice, LNP codelivery of the resulting melanoma PROTAV with a bi-adjuvant of oligonucleotide agonists for cyclic-GMP-AMP synthase (cGAS) and Toll-like receptor 9 (TLR9)^25,26^ resulted in potent multi-antigen-specific T cell responses. In a syngeneic melanoma mouse model, PROTAV, especially when combined with ICIs, not only promoted systemic antitumor immune responses, but also reduced tumor immunosuppression and promoted the tumor infiltration of antitumor T cells. In two melanoma syngeneic mouse models, relative to benchmark protein or peptide vaccines, PROTAV significantly promoted ICB therapeutic efficacy of tumors and increased the rates of tumor complete regression (CR). Remarkably, combining PROTAV and ICB achieved 100% eradication of large *Braf^V600E^* melanoma without recurrence in syngeneic mice. These results demonstrate the potential of PROTAVs for robust tumor combination immunotherapy.

**Fig. 1.**
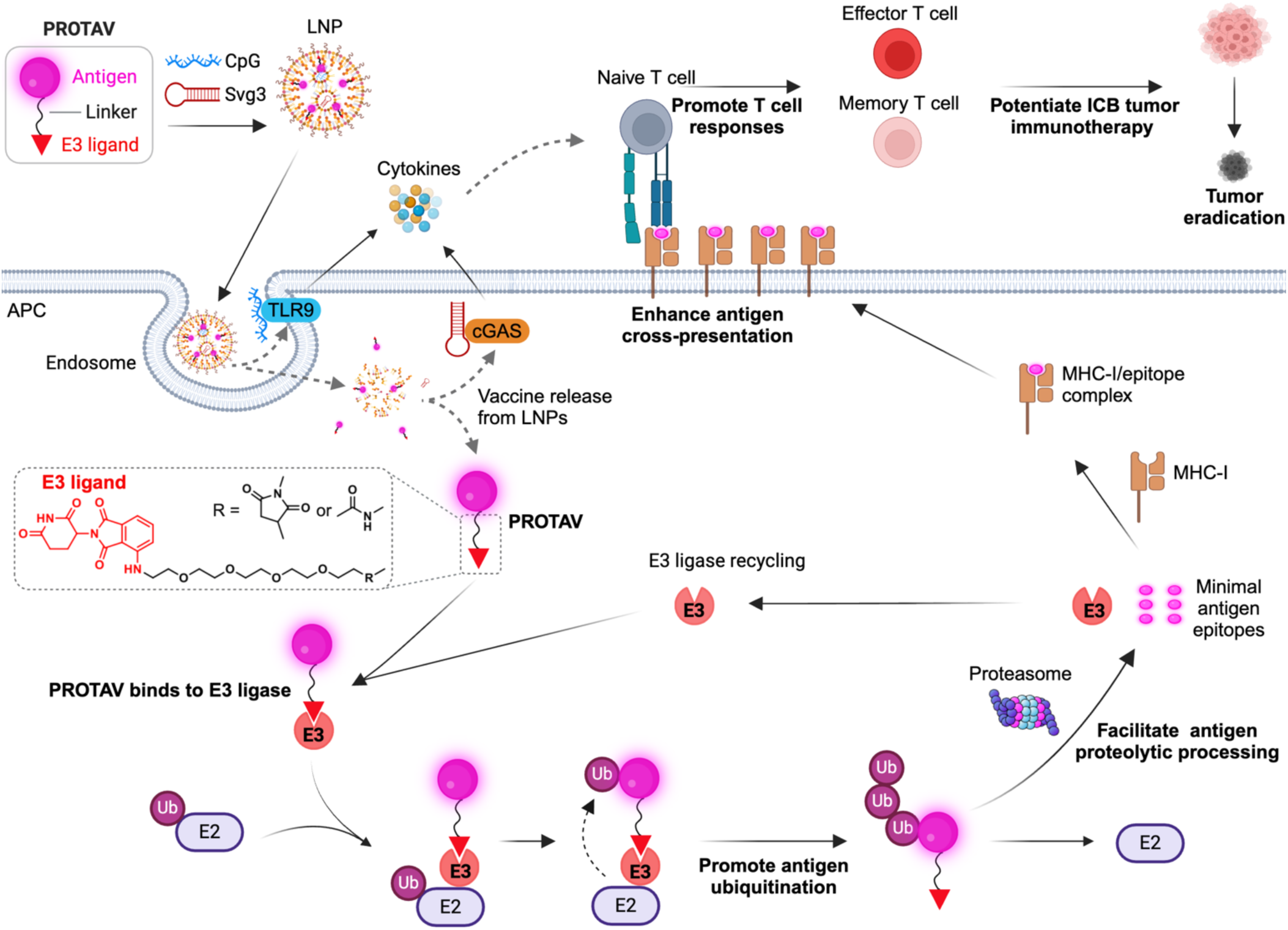
Schematic illustration of PROTAVs that facilitate antigen proteolytic processing in APCs and promote antigen cross-presentation to potentiate antitumor T cell responses for robust ICB combination immunotherapy of tumor. PROTAVs are covalent conjugates of a protein/peptide antigen and an E3 ligase-binding ligand (illustrated here: CRBN ligand pomalidomide), with a chemical linker in between. PROTAVs are broadly applicable for common types of tumor protein/peptide antigens, including proteins and synthetic long peptides. Upon delivery into APCs by LNPs, PROTAVs are bound to endogenous E3 ligases, which promoted the ubiquitination of the antigens in PROTAVs, facilitated antigen proteolytic processing by proteasome, promoted antigen cross-presentation to T cells, thereby potentiating antigen-specific T cell responses. A melanoma PROTAV was developed using a synthetic long peptide comprised of trivalent melanoma-associated antigens: Trp2, gp100, and Trp1 (TgT). The resulting PROTAV-TgT was co-loaded with biadjuvants, Svg3 and CpG, in LNPs for optimal PROTAV/adjuvant codelivery into APCs. The bi-adjuvants Svg3 and CpG are oligonucleotide immunostimulants that activate cGAS and TLR9, respectively, to elicit proinflammatory responses, including type-I interferon (IFN) responses. As a result, PROTAVs potentiated antitumor CD8^+^ cytotoxic T cell responses. In melanoma mouse models, the combination of PROTAV-TgT with ICB reduced immunosuppression and enhanced antitumor immunity in the tumor microenvironment. As a result, combining PROTAV-TgT with ICB facilitated tumor regression, and promoted the CR rates of tumors, including 100% eradication of large *Braf^V600E^* melanoma in syngeneic mice. LNP: lipid nanoparticle; cGAS: cyclic GMP-AMP synthase; Svg3: an oligonucleotide agonist for cGAS; TLR9: Toll-like receptor 9; CpG: an oligonucleotide agonist for TLR9; E2: E2 ligase; E3: E3 ligase; Ub: ubiquitin; APC: antigen-presenting cell; MHC: major histocompatibility complex; ICB: immune checkpoint blockade. Created in BioRender.

## Results

### Synthesis, linker optimization, and characterization of PROTAVs

A PROTAV is a modular molecular conjugate of a protein/peptide antigen, an E3 ligase-binding ligand^27^, and a linker in between (**Fig. 2a**). Depending on their amino acid sequences, tumor protein/peptide antigens have heterogeneous physicochemical properties and chemical reactivities. Thus, a broadly applicable, biocompatible, simple, and efficient method for PROTAC synthesis is highly desirable for developing PROTAVs for cancer immunotherapy. To this end, we tested PROTAV synthesis via two biocompatible and efficient conjugation chemistries: 1) N-hydroxysuccinimidyl (NHS) and 1-ethyl-3-(3-dimethylaminopropyl) carbodiimide (EDC) conjugation of NHS-modified E3 ligand with the primary amines of amino acids in antigens, and 2) thiol-maleimide conjugation of maleimide-modified E3 ligand with thiol on cysteine in antigens. We expect that the ubiquitous amino acids with primary amines (e.g., lysine) and thiol (i.e., cysteine) would enable broad application of these methods for PROTAV synthesis. Moreover, the modular structure of PROTAVs allows for the site-specific addition of reactive amino acids into antigens for optimal conjugation with E3 ligands. Various molecular ligands for many E3 ligase subtypes, such as cereblon (CRBN), have been studied in PROTACs, including multiple clinical studies of CRBN-based PROTACs.^28–32^ We chose pomalidomide, a commonly used CRBN ligand, as a representative E3 ligand in our PROTAV studies. Pomalidomide is an FDA-approved drug, which is expected to facilitate the clinical translation of pomalidomide-based PROTAV due to its clinically validated CRBN binding ability and the safety profile of pomalidomide.^30^ Note that, the structural modularity of PROTAVs allows for easy adjustment of E3 ligands.^27,33^ To synthesize PROTAVs, we first used chicken ovalbumin (OVA) as a model antigen, which can be proteolytically processed to generate a murine MHC-I-restricted minimal peptide epitope (SIINFEKL, OVA_257-264_) to elicit CD8^+^ T cell response.^34^

**Fig. 2.**
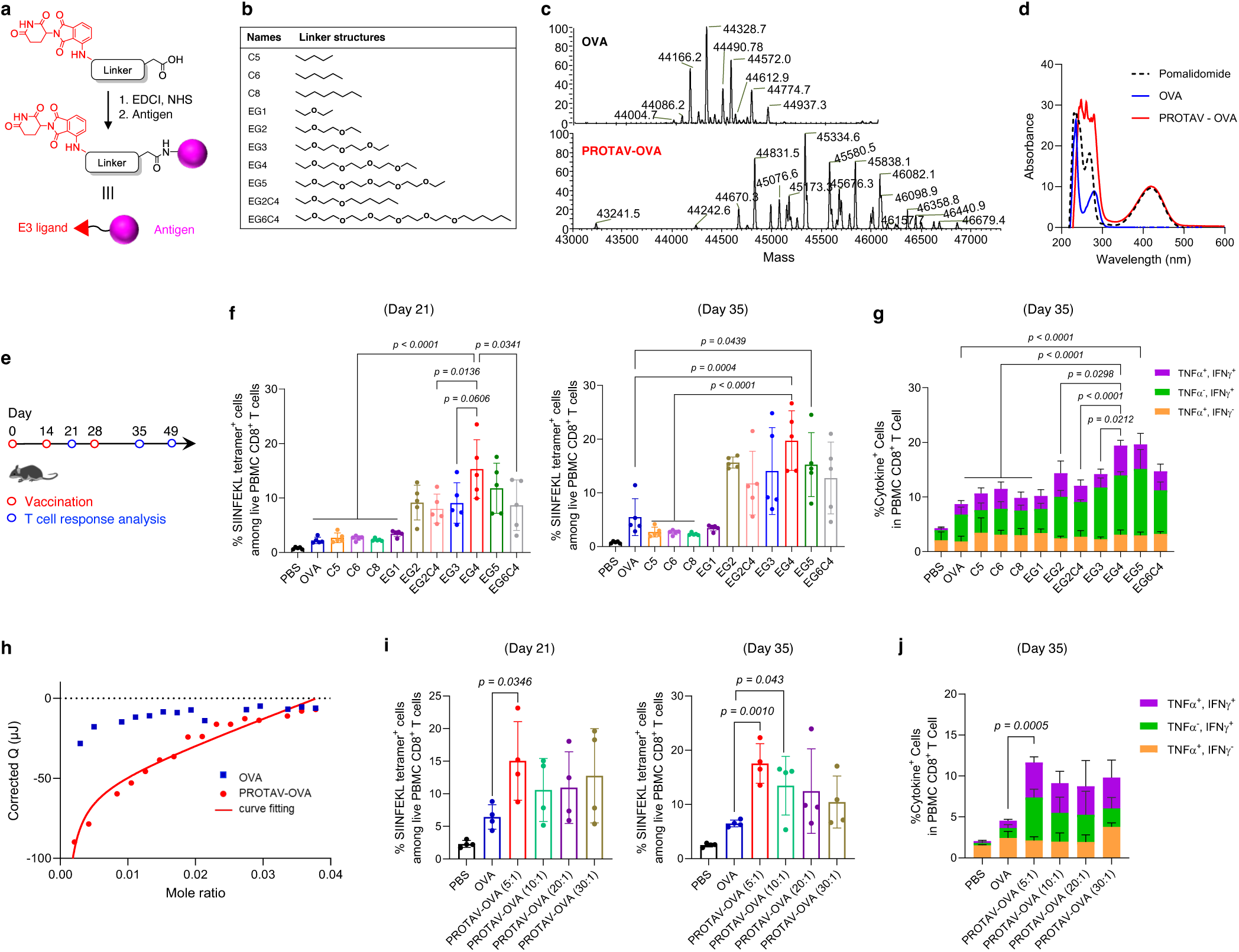
Linker optimization for PROTAV-OVA to elicit potent T cell response in mice. **a**, Representative synthesis route for PROTAVs via EDC-NHS conjugation. **b**, The chemical structures of a list of linkers screened for PROTAV-OVA. **c**, Deconvoluted ESI-MS spectra of OVA and PROTAV-OVA with an EG_4_ linker. **d**, UV-vis absorbance spectra of pomalidomide, OVA, and PROTAV-OVA (i.e., pomalidomide-OVA conjugates). The presence of pomalidomide-characteristic absorption (peak at 420 nm) in PROTAV-OVA suggests its successful synthesis. **e**, Timeline of T cell response study for PROTAV-OVA in mice. C57BL/6 mice (6-8 weeks; *n* = 5) were immunized with PROTAVs and controls, individually, by *s.c.* administration at tail base (day 0, day 14). CpG (2 nmole) adjuvant was mixed with OVA or PROTAV-OVA (10 μg). PBMCs were collected for T cell response analysis at indicated time points. **f, g**, H-2K^b^/SIINFEKL tetramer staining results (day 21, day 35) (**f**) and intracellular cytokine staining (day 35) (**g**) showing the fraction of SIINFEKL-specific T cell response among PBMC CD8^+^ T cells elicited by PROTAV-OVA candidates with various linkers. PROTAV-OVA with an EG_4_ linker consistently elicited one of the most potent T cell responses. **h**, Nano ITC results and fitted curve of the binding kinetics between PROTAV-OVA with CRBN protein, with OVA as a control. *K_d_* of PROTAV-OVA and CRBN binding was determined to be 1.2 μM using a one site-specific binding model. OVA and CRBN ITC data did not fit into the binding modeling, suggesting negligible binding. **i-j**, H-2K^b^/SIINFEKL tetramer staining results (day 21, day 35) (**i**) and intracellular cytokine staining (day 35) (**j**) showing the fraction of SIINFEKL-specific T cell response among PBMC CD8^+^ T cells elicited by PROTAV-OVA synthesized with four different feeding molar ratios of pomalidomide-EG_4_-NHS: OVA. Data represent mean ± s.e.m.; statistical analysis was conducted using one-way ANOVA with Bonferroni post-test.

Linker chemistry can be critical for molecular conjugate therapeutics^35^, including PROTACs^36^. We hypothesize that the linkers between E3 ligand and antigen in PROTAVs can be critical for its CRBN binding, antigen processing, and induction of T cell response. To screen for the optimal linker for PROTAV to elicit T cell response, we synthesized a series of PROTAVs with linkers of various water solubility (ethylene glycol or lipid) and length (**Fig. 2b**). PROTAV-OVA was first synthesized via NHS-EDC conjugation in two steps (**Fig. 2a**). First, we conjugated pomalidomide to an NHS-modified linker, as verified by electrospray ionization mass spectrometry (ESI-MS) (**Supplementary Fig. 1**). The resulting pomalidomide-linker showed great biostability, with limited degradation of pomalidomide-EG_4_ as an example after 2-hour incubation in serum, as measured by LC-MS (**Supplementary Fig. 2**). Second, we conjugated the resulting pomalidomide-linker-NHS with OVA via NHS conjugation with surface amines of OVA. The products were subject to ultracentrifugation purification. PROTAV-OVA synthesis was verified by 1) liquid chromatography-MS (LC-MS) (**Fig. 2c**), and 2) the presence of the characteristic UV-vis absorbance of pomalidomide (peak absorbance at 420 nm) in PROTAV-OVA (**Fig. 2d**). Conveniently, the linear correlation of pomalidomide absorption with PROTAV-OVA concentrations allows for easy estimation of the average pomalidomide copy numbers per PROTAV-OVA via UV-vis absorbance (**Supplementary Fig. 3a**). Next, by subcutaneous (*s.c.*) injection at tail base (day 0 and day 14), C57BL/6 mice were immunized with these PROTAV candidates individually, with unmodified OVA and PBS as controls and immunostimulant CpG-1826 oligonucleotide (a TLR9 agonist) (**Supplementary Table 1**) as adjuvant (**Fig. 2e**). H-2K^b^/SIINFEKL tetramer staining (day 21, day 35) showed that, while all PROTAV-OVA(s) promoted the fraction of SIINFEKL-specific CD8^+^ T cells among peripheral blood mononuclear cell (PBMC) CD8^+^ T cells, PROTAV-OVA with a tetrameric ethylene glycol (EG_4_) linker consistently elicited the highest T cell responses (**Fig. 2f**). Further, as shown by *ex vivo* antigen restimulation of PBMC T cells followed by intracellular cytokine staining (day 35), PROTAV-OVA with an EG_4_ linker elicited the most multifunctional CD8^+^ T cells that produced interferon-γ (IFN-γ) and tumor necrosis factor-α (TNF-α) (**Fig. 2g**). These results demonstrated that PROTAV-OVA with an EG_4_ linker elicited a large quantity and a good quality of T cell responses. This, together with its relatively short length, allows EG_4_ linker to be selected for further studies. Moreover, physical mixture of OVA and pomalidomide elicited comparable levels of SIINFEKL-specific CD8^+^ T cell response relative to OVA alone (**Supplementary Fig. 4**). This not only verifies the benefit of covalent pomalidomide-antigen conjugation for T cell response, but also rules out the impact from the intrinsic immunomodulatory effect of pomalidomide on the ability of PROTAV to elicit T cell responses. In a molecular binding assay using nano isothermal titration calorimeter (ITC), PROTAV-OVA showed strong binding affinity with CRBN (dissociation constant *K_d_* = 1.2 μM), in contrast to negligible binding between OVA and CRBN (**Fig. 2h**, **Supplementary Fig. 5**). Relative to pomalidomide binding to CRBN (*K_d_* = 156 nM)^37^, the reduced binding affinity between CRBN and pomalidomide in PROTAV-OVA is presumably caused by the structural hindrance of OVA for CRBN binding. Nonetheless, these results provide the basis for avid CRBN binding in APCs to facilitate antigen ubiquitination and proteolytic processing.

The copy number of E3 ligands per antigen can also be critical for their E3 ligase binding and antigen processing. To test PROTAV-OVA with varying pomalidomide copy numbers per antigen, we synthesized PROTAV-OVA with different pomalidomide-EG_4_-NHS: OVA feeding molar ratios at 5: 1, 10: 1, 20: 1, and 30: 1, resulting in PROTAV-OVA with averages of 2.1, 6.3, 8.0, and 9.2 pomalidomide copies per OVA, respectively, as measured using a method of free-amine-reactive fluorescent labeling (**Supplementary Fig. 3b**). To study their T cell responses, mice were immunized as above with CpG-adjuvanted PROTAV-OVA with different pomalidomide copies, respectively (**Fig. 2e**). H-2K^b^/SIINFEKL tetramer staining revealed that a pomalidomide: OVA feeding ratio of 5:1 elicited one of the highest fractions of SIINFEKL-specific PBMC CD8^+^ T cells (**Fig. 2i**). Consistently, PROTAV-OVA with pomalidomide: OVA feeding ratio of 5:1 also resulted in the most fraction of multifunctional cytokine-producing PBMC CD8^+^ T cells upon antigen restimulation (**Fig. 2j**). This, together with its simplicity of synthesis and quality control, allow us to choose this feeding ratio for further studies. Next, we compared NHS-EDC conjugation with thiol-maleimide conjugation for PROTAV synthesis, which conjugate pomalidomide with the intrinsic primary amine and thiol on antigens, respectively. UV-vis absorbance confirmed that comparable copy numbers of pomalidomides were grafted per OVA by the two methods (**Supplementary Fig. 3c**). After immunization of mice with the resulting PROTAV-OVA, H-2K^b^/SIINFEKL tetramer staining confirmed that PROTAV-OVA synthesized using these two methods elicited comparable fractions of SIINFEKL-specific PBMC CD8^+^ T cells (**Supplementary Fig. 6**).

### PROTAVs facilitated antigen proteolysis and promoted its cross-presentation in DCs *in vitro* and *in vivo*

The proteolytic processing of protein/peptide antigens by 26S proteasome is a critical step in the antigen processing cascade prior to presentation of the resulting minimal antigen epitopes from APCs to T cells. We hypothesize that, by directly conjugating E3 ligands on antigens, PROTAVs enhance antigen ubiquitination, and thus expedite antigen processing and presentation, resulting in promoted T cell activation. We studied antigen proteolytic processing (*i.e.*, degradation) of PROTAVs using PROTAV-OVA prepared from a fluorogenic DQ-OVA conjugate. The DQ fluorescence of DQ-OVA is self-quenched until OVA degradation^38^, allowing for a semi-quantitative study of OVA degradation. After treatment of DC2.4 cells with PROTAV-DQ-OVA *vs*. DQ-OVA for a series of durations up to 72 h, flow cytometry showed elevated DQ fluorescence of cells treated with PROTAV-DQ-OVA relative to DQ-OVA at the corresponding time points (**Fig. 3a-b**). We further studied this by confocal microscopy. After treatment of DCs as above for only 0.5 h, PROTAV-DQ-OVA started to show enhanced DQ fluorescence recovery relative to DQ-OVA (**Fig. 3c**). Following a 0.5-h treatment, cells were washed to remove extracellular PROTAV-OVA-DQ and then incubated for an additional 5 h. Confocal microscopy showed that, compared to the initial 0.5-h treatment, the ratio of cytosolic to endolysosomal DQ fluorescence in DCs increased after the extended incubation (**Fig. 3d**). This not only suggests enhanced antigen degradation by PROTAV at the subcellular levels, but also demonstrated effective delivery of PROTAV-OVA to the cytosol, which is required for antigens to bind to E3 ligase and undergo proteolytic processing. At the molecular level, western blot (WB) demonstrated that PROTAV-OVA facilitated OVA degradation relative to OVA in DCs (**Fig. 3e, Supplementary Fig. 7a, b**). Mechanistically, co-immunoprecipitation (co-IP) showed that PROTAV enhanced the fraction of OVA and ubiquitin colocalization in DCs, suggesting that PROTAV promoted OVA ubiquitination (**Fig. 3f, Supplementary Fig. 7c**). These results suggest that PROTAV facilitated antigen proteolytic processing, a prerequisite for antigen presentation.

**Fig. 3.**
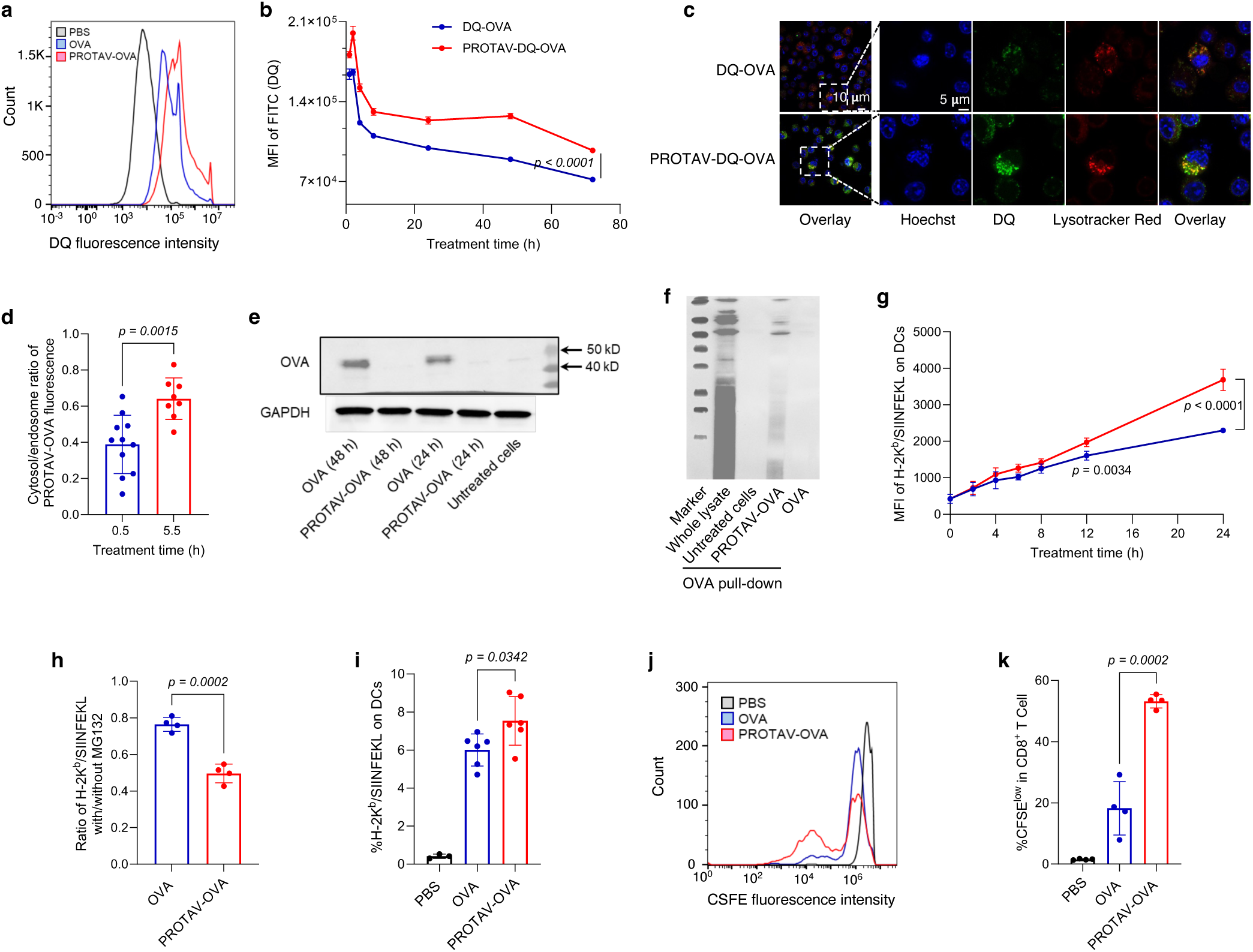
PROTAV promoted antigen proteolytic processing and facilitated antigen presentation in DCs *in vitro* and *in vivo*. **a-b**, Representative flow cytometry graph (**a**) and MFI quantified from flow cytometry data (**b**) showing that PROTAV-OVA-DQ facilitated OVA-DQ degradation (antigen processing) than OVA-DQ in DCs after a 24-h treatment (**a**) and over the course of 72 h post treatment (**b**). **c**, Confocal microscopy images showing stronger DQ fluorescence recovery of DCs treated with PROTAV-OVA-DQ than cells treated with OVA-DQ (1 h). **d**, The ratios of cytosolic over endolysosomal DQ fluorescence intensities in DCs, after treatment for 0.5 h and 5.5 h, respectively. Cells were treated with PROTAV-OVA-DQ for 0.5 h. One group of cells were subject to imaging, and the other group of cells were washed with cell culture medium to remove extracellular PROTAV-OVA-DQ, followed by another 5-h incubation before imaging. Data were quantified from confocal microscopy images. **e**, A WB image showing that after DCs were treated with PROTAV-OVA and OVA, respectively, PROTAV-OVA resulted in elevated OVA degradation relative to OVA after the corresponding treatment durations (24 h or 48 h). **f**, A co-IP image showing that PROTAV-OVA promoted the ubiquitination of OVA relative to unconjugated OVA in DC2.4 cells after treatment for 12 h. **g**, MFI of H-2K^b^/SIINFEKL on DCs quantified from flow cytometry data showing that PROTAV-OVA promoted the presentation of SIINFEKL antigen epitope on DCs over the course of 24-h treatment. H-2K^b^/SIINFEKL on DCs was stained using a dye-labeled antibody. **h**, Protease inhibitor MG132 offsets the ability of PROTAV-OVA to facilitate antigen presentation on DCs. In **g-h**, vaccines were transfected into DCs using lipofectamine 2000. **i**, Quantified flow cytometry results showing the percentage of H-2K^b^/SIINFEKL-positive DCs among total live DCs from draining inguinal lymph nodes in C57BL/6 mice treated with PROTAV-OVA-DQ or OVA-DQ, respectively. Vaccines were *s.c.* administered at mouse tail base (dose: 10 μg OVA or PROTAV-OVA, 2 nmole CpG). **j**-**k**, Representative flow cytometry graphs (**j**) and quantified flow cytometry results showing the CFSE-low OT-1 CD8^+^ T cells upon co-culture (96 h) with DCs pre-treated (24 h) with PROTAV-OVA or OVA, respectively. This suggests that relative to OVA, PROTAV-OVA promoted the ability of DCs to expand antigen-specific T cells, as indicated by the CFSE signal dilution of OT-1 cells during cell division. 10 mg/mL PROTAV-OVA or OVA was used in all *in vitro* studies. Data represent mean ± s.e.m.; statistical analysis was conducted by one-way ANOVA with Bonferroni post-test.

To study antigen presentation, C57BL/6 mouse-derived bone marrow-derived DCs (BMDCs) were treated with PROTAV-OVA *vs*. OVA for a series of durations of up to 24 h, followed by antibody staining and flow cytometric analysis of the complex of SIINFEKL and H-2K^b^ (an MHC-I subtype). CpG-1826 oligonucleotide, an agonist for murine TLR9,^26^ was used as an immunostimulant adjuvant in the above vaccine. As a result, after treatment for 12 - 24 h, relative to OVA, PROTAV-OVA enhanced the level of H-2K^b^/SIINFEKL on DCs, suggesting that PROTAV promoted antigen presentation (**Fig. 3g**). Note that, relative to OVA, PROTAV-OVA did not significantly enhance SIINFEKL presentation until 12 h after treatment. This is likely because the cytosolic delivery of OVA or PROTAV-OVA is time-consuming and inefficient, involving endocytosis and endosome escape; and shortly after treatment, pre-existing E3 ligases suffice to efficiently process both OVA and PROTAV-OVA that have reached the cytosol. Further, proteasome inhibitor (MG132)^39^ reduced the level of SIINFEKL presentation on DCs treated with PROTAV-OVA or OVA as above (**Fig. 3h**), verifying that PROTAV enhances antigen presentation in a proteasome-dependent manner. To study this *in vivo*, C57BL/6 mice were *s.c.* injected at tail base with PROTAV-OVA *vs*. OVA, with CpG adjuvant. After up to 4 days, analysis of draining inguinal lymph nodes revealed that PROTAV-OVA elevated SIINFEKL presentation levels on intranodal DCs throughout this course (**Fig. 3i**). To further study PROTAV-treated DCs for antigen presentation to T cells, BMDCs were treated as above with CpG-adjuvanted PROTAV-OVA *vs.* OVA for 24 h. SIINFEKL-specific OT-1 CD8^+^ T cells with engineered SIINFEKL-specific T cell receptors were isolated from OT-1 transgenic mice and stained with carboxyfluorescein succinimidyl ester (CFSE). The above vaccine-treated BMDCs were then cocultured with CFSE-stained OT-1 CD8^+^ T cells for 96 h, followed by flow cytometric analysis of CFSE fluorescence signal intensities of OT-1 T cells to determine cell division. As a result, OT-1 CD8^+^ T cells cocultured with PROTAV-OVA-treated DCs underwent significantly more CFSE signal dilution and cell division (**Fig. 3j-k**). This suggests that PROTAV-OVA enhanced antigen presentation from DCs to promote T cell activation. For further mechanistic understanding, we employed RNA sequencing (RNA-seq) for transcriptomic analysis of BMDCs treated with CpG-adjuvanted PROTAV-OVA or OVA, respectively. Gene ontology (GO) enrichment analysis revealed that, relative to OVA, PROTAV-OVA significantly activated the ubiquitin and immune response-related pathways (**Supplementary Fig. 8**). Taken together, these data demonstrated that PROTAV promoted proteasome-dependent antigen proteolytic processing, facilitated antigen presentation, and enhanced T cell activation, providing the basis for PROTAV to promote T cell responses.

### Multivalent melanoma PROTAVs elicited potent antigen-specific T cell responses

Depending on the amino acid sequences, protein or peptide antigens have highly heterogeneous tertiary structures and structurally accessible amino acids for E3 ligand conjugation. Synthetic long peptides, which cover the minimal peptide epitopes to elicit antitumor T cell responses, are extensively studied in cancer therapeutic vaccines. Relative to minimal peptide epitopes, synthetic long peptide antigens allow for de novo antigen processing and complexation with MHC in APCs for potent T cell activation with minimal tolerance.^40,41^ To explore the applicability of PROTAVs in synthetic long peptide vaccines, we studied PROTAV for trivalent melanoma-associated antigens. Melanoma represents the most prevalent skin cancer, and advanced melanoma has a <30% 5-year survival rate. Though ICB^3,5,42^ has improved melanoma treatment outcomes, most melanoma patients do not respond to ICB^43,44^ or suffer from irAEs.^3–5^ Cancer therapeutic vaccines can improve ICB therapeutic efficacy by inducing antitumor immune cells as ICB targets, upregulating immune checkpoints, and reducing tumor immunosuppression.^8,45–48^ Synthetic long peptides have shown the benefit to elicit T cell responses with minimal immune exhaustion,^49^ a process that is hinged to peptide proteolytic processing in order for antigen presentation. To study PROTAV for melanoma, we designed a trivalent antigenic fusion peptide, TgT, for three long peptides of MHC-I melanoma TAAs: **T**rp2_175-194_, **g**p100_16-35_, and **T**rp1_450-469_ (**Fig. 4a**).^50–53^ We expect TgT to elicit multi-specific CD8^+^ T cells to overcome tumor antigen heterogeneity and the associated immune escape for optimal melanoma therapeutic efficacy.^40,54^ Moreover, multiple N-/C-terminal K and C were added to promote pomalidomide conjugation via maleimide-thiol conjugation and ubiquitination on lysine, respectively. As predicted by AlphaFold3,^55^ cysteines in the TgT peptide have overall good spatial accessibility, which is expected to allow for their chemical conjugation with maleimide-EG_4_-pomlidomide (**Fig. 4b**). Thus, under a reductive condition to minimize dithiol formation from intrinsic cysteines, we synthesized PROTAV-TgT by conjugating TgT with maleimide-EG_4_-pomalidomide, as verified by LC-MS (**Supplementary Fig. 9**).

**Fig. 4.**
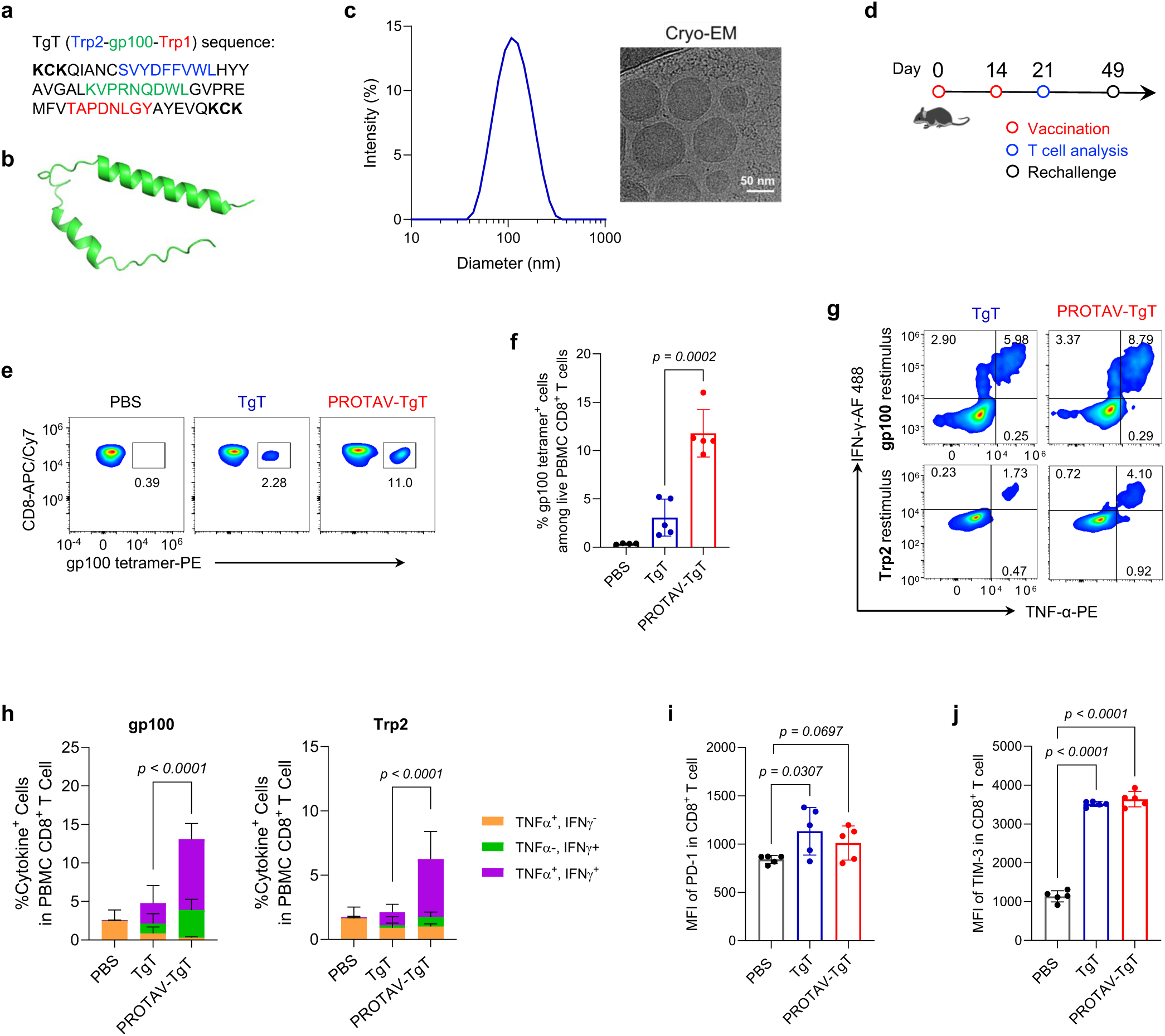
Multivalent PROTAV-TgT elicited potent antigen-specific T cell responses. **a**, Sequence of the TgT multivalent antigenic peptide used in PROTAV-TgT. Minimal antigenic epitopes were underlined. Multiple terminal K and C were added to promote pomalidomide conjugation via maleimide-thiol conjugation and ubiquitination on lysine, respectively. **b**, Structure of TgT predicted by AlphFold3. **c**, DLS data and a cryo-EM image showing the hydrodynamic sizes and morphology of SM-102 LNPs co-loaded with PROTAV-TgT, CpG, and Svg3. CpG: Svg3 molar ratio was 2: 1. Nucleic acid phosphate over lipid nitrogen N:P ratio was 6:1. **d**, Timeline of PROTAV-TgT immunization study in mice. C57BL/6 mice (6-8 weeks) were immunized with PROTAV-TgT and TgT vaccines, respectively (*s.c.* at tail base, day 0 and day 14). Dose: 20 ug antigen, 2 nmole CpG, and 1 nmole Svg3. **e-j**, PROTAV-TgT elicited potent multivalent T cell responses while upregulating immune checkpoint levels on T cells in mice. **e-f**, Representative flow cytometry graphs (**e**) and quantified flow cytometry data for tetramer staining results showing that relative to TgT, PROTAV-TgT promoted gp100-specific CD8^+^ T cell responses (day 21). **g-h**, Representative flow cytometry graphs (**g**) and quantified intracellular cytokine staining results (**h**) of CD8^+^ T cells upon ex vivo restimulation of PBMC CD8^+^ T cells with gp100 and Trp2 peptides, respectively, suggesting that relative to TgT, PROTAV-TgT promoted multifunctional T cell responses producing antitumor cytokines. **i-j**, MFI of PD-1 (**i**) and Tim-3 (**j**) on total live PBMC CD8^+^ T cells in the as-immunized mice, suggesting that PROTAV-TgT upregulated these immune checkpoint levels on T cells. Data represent mean ± s.e.m. (*n* = 5); statistical analysis was conducted using one-way ANOVA with Bonferroni post-test unless denoted otherwise.

Peptides and their derivatives, such as TgT and PROTAV-TgT, often have low water solubility and poor delivery efficiency to lymphoid tissues, such as draining lymph nodes where APCs process and present antigens to T cells. Moreover, peptide vaccines are overall poorly immunogenic, which demands immunostimulant adjuvants to elicit T cell responses. Yet, many of current molecular adjuvants, including TLR agonists such as TLR9 agonist CpG, still elicit suboptimal T cell responses in human when used together with peptide antigens. This is in part because of the restricted TLR9 expression in limited subsets of APCs (e.g., plasmacytoid DCs) and their limited ability to elicit type-I IFN response, which is critical to elicit T cell response. Lastly, a physical mixture of peptide antigens and adjuvants often randomly disseminates with poor co-delivery to lymphoid APCs, causing suboptimal T cell responses. To address these issues, first, we recently developed an oligonucleotide-based cGAS agonist, Svg3.^25^ Svg3 selectively activates murine and human cGAS, which is ubiquitously expressed in various APC subsets, to elicit strong type-I IFN responses. Thus, we combine Svg3 and CpG as bi-adjuvant for PROTAV to concurrently activate cGAS and TLR9, thereby eliciting potent T cell responses. Second, we used ionizable SM-102 LNPs, which have been used in a COVID-19 mRNA vaccine, as nanocarriers to co-load PROTAV-TgT and adjuvants Svg3 and CpG. PROTAV-TgT/Svg3/CpG-co-loaded LNPs were synthesized via rapid microfluidic mixing, with the Svg3: CpG molar ratio of 1:2 and an overall N:P ratio of 6:1. The resulting PROTAV-TgT/Svg3/CpG-loaded LNPs displayed hydrodynamic diameters of 157 ± 5 nm (mean ± SD) with great homogeneity (PDI: 0.128), as measured by hydrodynamic light scattering (DLS) (**Fig. 4c**), which is expected to allow for efficient delivery into draining lymph nodes. Cryogenic electron microscopy (cryo-EM) further revealed the exterior and internal morphology of PROTAV-TgT/Svg3/CpG-loaded LNPs (**Fig. 4c**).

Next, we studied the resulting PROTAV-TgT vaccines (PROTAV-TgT/Svg3/CpG-loaded LNPs) to elicit T cell responses. C57BL/6 mice were immunized with PROTAV-TgT vaccine and TgT vaccine (TgT/Svg3/CpG-loaded LNPs), respectively (*s.c*. at tail base, day 0 and day 14) (**Fig. 4d**). On day 21, PBMC T cells were analysed by flow cytometry to assess TgT-specific T cell responses. Specifically, gp100/H-2D^b^ tetramer staining showed that relative to TgT, PROTAV-TgT promoted gp100-specific CD8^+^ T cell responses (**Fig. 4e, f**). Moreover, intracellular cytokine staining revealed that PROTAV-TgT promoted multifunctional CD8^+^ cell responses (**Fig. 4g, h**). Further, PROTAV-TgT vaccines upregulated the expression levels of immune checkpoints programmed death receptor 1 (PD-1) and T-cell immunoglobulin and mucin domain 3 (Tim-3) on PBMC CD8^+^ T cells (**Fig. 4i, j**), presumably due to the innate and adaptive immunostimulation upon immunization as well as enhanced antigen exposure for T cells, especially for PROTAV. This suggests that PROTAV sensitized immune checkpoints for ICB therapy, providing the basis to combine ICB (e.g., PD-1 antibodies or αPD-1) with PROTAV for optimal tumor therapeutic efficacy. Lastly, flow cytometry analysis of immune memory-associated phenotypical markers CD44 and CD62L on PBMC CD8^+^ T cells demonstrates that PROTAV-TgT elicited great CD8^+^ T cell memory relative to TgT (**Supplementary Fig. 10**), which is critical for long-term tumor therapeutic efficacy with minimal tumor relapse.

### PROTAV-TgT, in combination with ICIs, reduced immunosuppression in tumor microenvironment (TME)

TME is primarily where tumor cells suppress antitumor immunity.^56^ In addition to systemic antitumor immunomodulation, modulating the tumor immune milieu is pivotal to reducing immunosuppression and promoting tumor immunotherapeutic efficacy.^57^ Using a B16F10 melanoma mouse model, we studied the systemic and TME immunomodulation by PROTAV-TgT, alone or combined with ICIs. Tumors were *s.c.* inoculated on mouse flank. Treatment was initiated when tumor sizes reached *ca*. 50 mm^3^. Specifically, mice were immunized with PROTAV-TgT, with PBS or TgT as controls by *s.c*. administration at tail base that allows for lymphatic draining to immunomodulatory lymph nodes, and mice were treated with αPD-1 by intraperitoneal (*i.p.*) administration. After two doses of vaccines and three doses of αPD-1, tumors, lymph nodes, along with PBMCs were collected for immune analysis (**Fig. 5a**). We first analyzed gp100-specific CD8^+^ T cells in tumors by gp100/H-2D^b^ tetramer staining. Relative to TgT, PROTAV-TgT, alone or combined with ICIs, respectively, promoted the tumor infiltration of gp100-specific CD8^+^ T cells (**Fig. 5b**). Importantly, relative to TgT + αPD-1, PROTAV-TgT + αPD-1 promoted the fraction of granzyme B-producing CD8^+^ T cells among total CD8^+^ T cells in melanoma (**Fig. 5c**). These data suggest that PROTAV-TgT + αPD-1 promoted the quantity and quality of tumor-infiltrating anti-tumor CD8^+^ T cells. Moreover, the ratio of CD8^+^ T cells over regulatory T cells (Treg) (CD8^+^/Treg) in tumor, which is predictive for effective tumor immunotherapy, was enhanced by PROTAV-TgT + αPD-1 (**Supplementary Fig. 11**). Importantly, relative to PROTAV-TgT or αPD-1 alone, PROTAV-TgT + αPD-1 reduced the fractions of PD-1^+^ intratumoral CD8^+^ T cells and the PD-1 expression level on these cells, and reduced the number of intratumoral terminally exhausted PD-1^+^Tim-3^+^ CD8^+^ T cells that are featured with reduced proliferative capacity, diminished antitumor cytokine production, and impaired antitumor cytotoxic activity (**Fig. 5d-e**). Collectively, these results suggested that PROTAV-TgT + αPD-1 promoted the antitumor cell density and functionality in TME. We then studied the systemic immunomodulation in peripheral blood and lymph nodes in the above treated mice. Again, gp100/H-2D^b^ tetramer staining of PBMCs showed that, relative to TgT, PROTAV-TgT, alone or combined with αPD-1, elevated the fraction of gp100-specific CD8^+^ T cells among PBMC CD8^+^ T cells (**Fig. 5f**). Similarly, relative to TgT, PROTAV-TgT, alone or combined with αPD-1, promoted the lymph node infiltration of gp100-specific CD8^+^ T cells (**Fig. 5g**). Further, PROTAV-TgT promoted the activation of type 1 conventional DCs (cDC1) in lymph nodes (**Fig. 5h**). cDC1 are DC subsets specialized in modulating cellular immune responses against cancer.^58^ These results suggest that PROTAV-TgT efficiently activated cDC1, which subsequently elicit potent TgT-specific CD8^+^ T cell responses. Overall, these results demonstrate that PROTAV-TgT, especially when combined with ICIs, increased antitumor immunity and reduced TME immunosuppression and enhanced TME antitumor immunity, while activating systemic antitumor T cell responses in tumor-bearing mice.

**Fig. 5.**
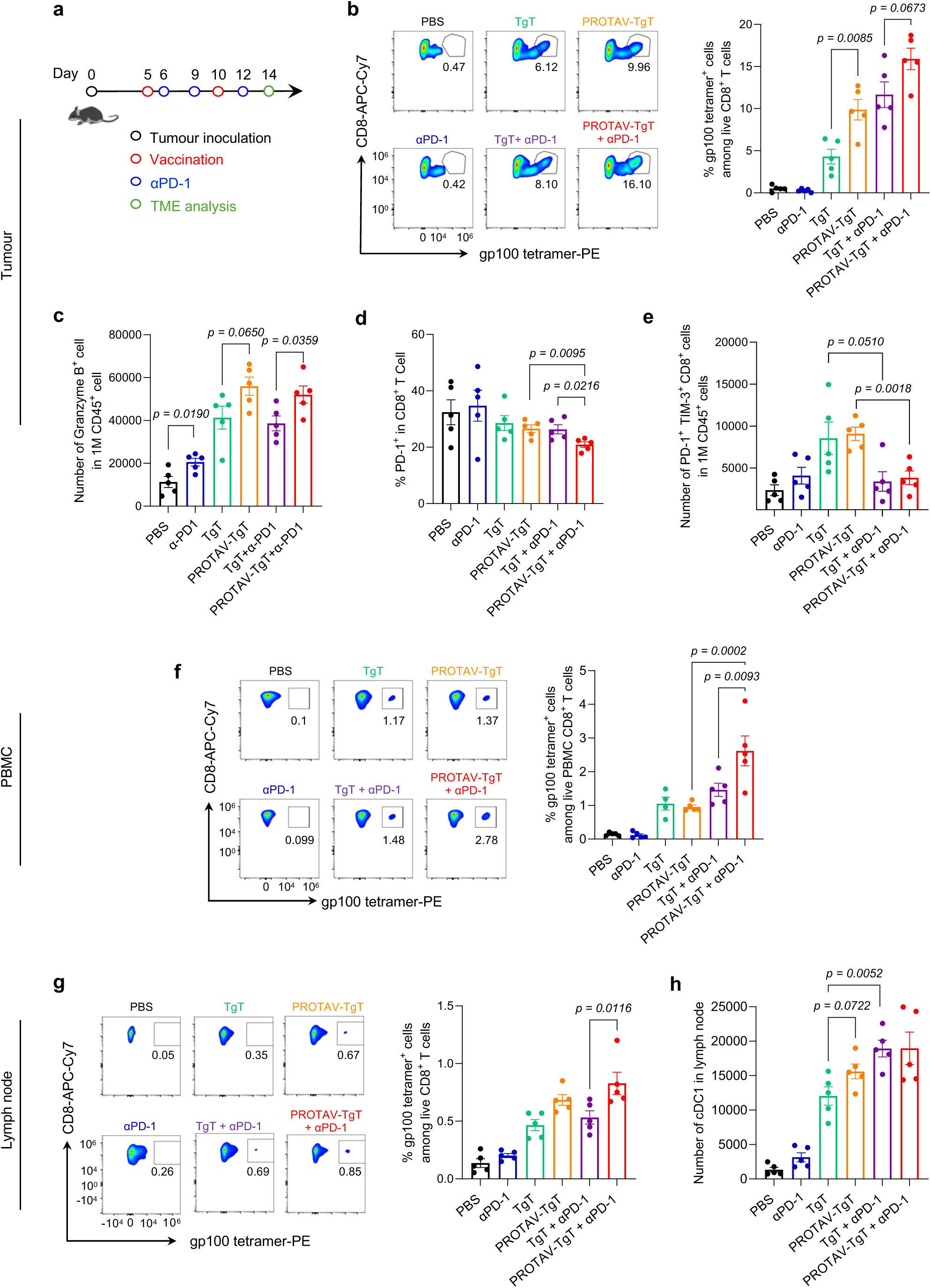
PROTAV-TgT, in combination with αPD-1, reduced immunosuppression in tumor microenvironment. **a** Timeline for the study of systemic and TME immunomodulation by PROTAV-TgT, alone or combined with αPD-1, in C57BL/6 mice with B16F10 melanoma. **b**, Representative flow cytometry graphs (left) and quantified flow cytometry results (right) of gp100/H-2D^b^ tetramer staining of intratumoral gp100-specific CD8^+^ T cells. Relative to TgT, PROTAV-TgT, alone or combined with αPD-1, promoted the tumor infiltration of antitumor CD8^+^ T cells, as shown by gp100-specific CD8^+^ T cells analyzed as an example by tetramer staining. **c**, Flow cytometry results showing the fractions of granzyme B-producing CD8^+^ T cells among total live CD8^+^ T cells from as-treated tumors. Relative to TgT + aPD-1, PROTAV-TgT + aPD-1 promoted the fraction of granzyme B-producing CD8^+^ T cells among total CD8^+^ T cells in melanoma. **d-e** Flow cytometry results showing the fractions of PD-1 expression on the CD8^+^ T cells (**d**) and the number of PD-1^+^ TIM-3^+^ CD8^+^ T cells in 1M CD45^+^ cell (**e**), **f**, Representative flow cytometry graphs (left) and quantified flow cytometry results (right) of gp100/H-2D^b^ tetramer staining of PBMC gp100-specific CD8^+^ T cells from mice treated as above. Relative to TgT, PROTAV-TgT, alone or combined with αPD-1, promoted the fraction of gp100-specific CD8^+^ T cells among total live PBMC CD8^+^ T cells. **g**, Representative flow cytometry graphs (left) and quantified flow cytometry results (right) of gp100/H-2Db tetramer staining of gp100-specific CD8^+^ T cells among lymph node total live CD8^+^ T cells. **h**, Quantified flow cytometry results showing the numbers of cDC1 in lymph nodes. Relative to TgT, PROTAV-TgT, alone or combined with αPD-1, promoted the lymph node infiltration of gp100-specific CD8^+^ T cells (**g**) and cDC1 cells (**h**). Data represents s.e.m. (*n* = 5); statistical analysis was conducted using one-way ANOVA with Bonferroni post-test.

### Combining PROTAV-TgT with ICIs promoted melanoma CR rates and eradicated 100% of *Braf^V600E^* melanoma in mice

Encouraged by the ability of PROTAV-TgT for TME and systemic antitumor immunomodulation in tumor-bearing mice, we then assessed PROTAV-TgT, alone or combined with ICB (i.e., αPD-1), for melanoma therapy. We first studied this in a syngeneic mouse model of *s.c*. B16F10 melanoma. Mice with established tumors (initial tumor size: ca. 40 mm^3^) were treated with: 1) PROTAV-TgT, with PBS or TgT as controls (*s.c*. immunization at tail base, days 3, 9, 15, 21), and 2) αPD-1 (*i.p.* administration, day 5, 8, 11, 14, 17, 20, 23) (**Fig. 6a**). Again, PROTAV-TgT or TgT were co-loaded with CpG and Svg3 in SM-102 LNPs. Because ICB prior to vaccines induces dysfunctional subprimed T cells and renders tumor resistant to immunotherapy,^59^ ICB was initiated one day post priming immunization to minimize the otherwise antitumor T cell immune dysfunctionality. Tumor growth showed that, relative to TgT, PROTAV-TgT alone already showed slightly stronger tumor growth (**Fig. 6b-d**). Interestingly, when combined with ICIs, PROTAV-TgT significantly facilitated the tumor regression and promoted the therapeutic response of B16F10 melanoma in syngeneic mice. Consistently, Kaplan-Meier mouse survival curves showed that PROTAV-TgT + αPD-1 resulted in significant survival benefits over controls such as PROTAV-TgT or ICIs alone, or TgT + αPD-1 combination. PROTAV-TgT + αPD-1 resulted in 4/8 tumor CR and long-term mouse survival without recurrence, in contrast to 1/8 for PROTAV-TgT and none for the other treatments (**Fig. 6c-d**). The median survival of mice treated with PROTAV-TgT + αPD-1 reached 58 days, a two-fold increase over TgT + αPD-1, PROTAV-TgT, or αPD-1. Moreover, bi-adjuvant Svg3/CpG outperformed single adjuvant CpG for PROTAV-TgT + αPD-1 to inhibit melanoma progression and enhance the rate of melanoma CR, demonstrating the benefits of biadjuvant for optimal tumor therapeutic efficacy (**Supplementary Fig 12**). Lastly, during the treatment course, PROTAV-TgT + αPD-1 did not cause any significant mouse body weight changes relative to control treatments (**Supplementary Fig. 13**), suggesting the promising safety of this treatment strategy. Collectively, these results clearly demonstrated that PROTAV-TgT promoted the ICB combination therapeutic efficacy of melanoma.

**Fig. 6.**
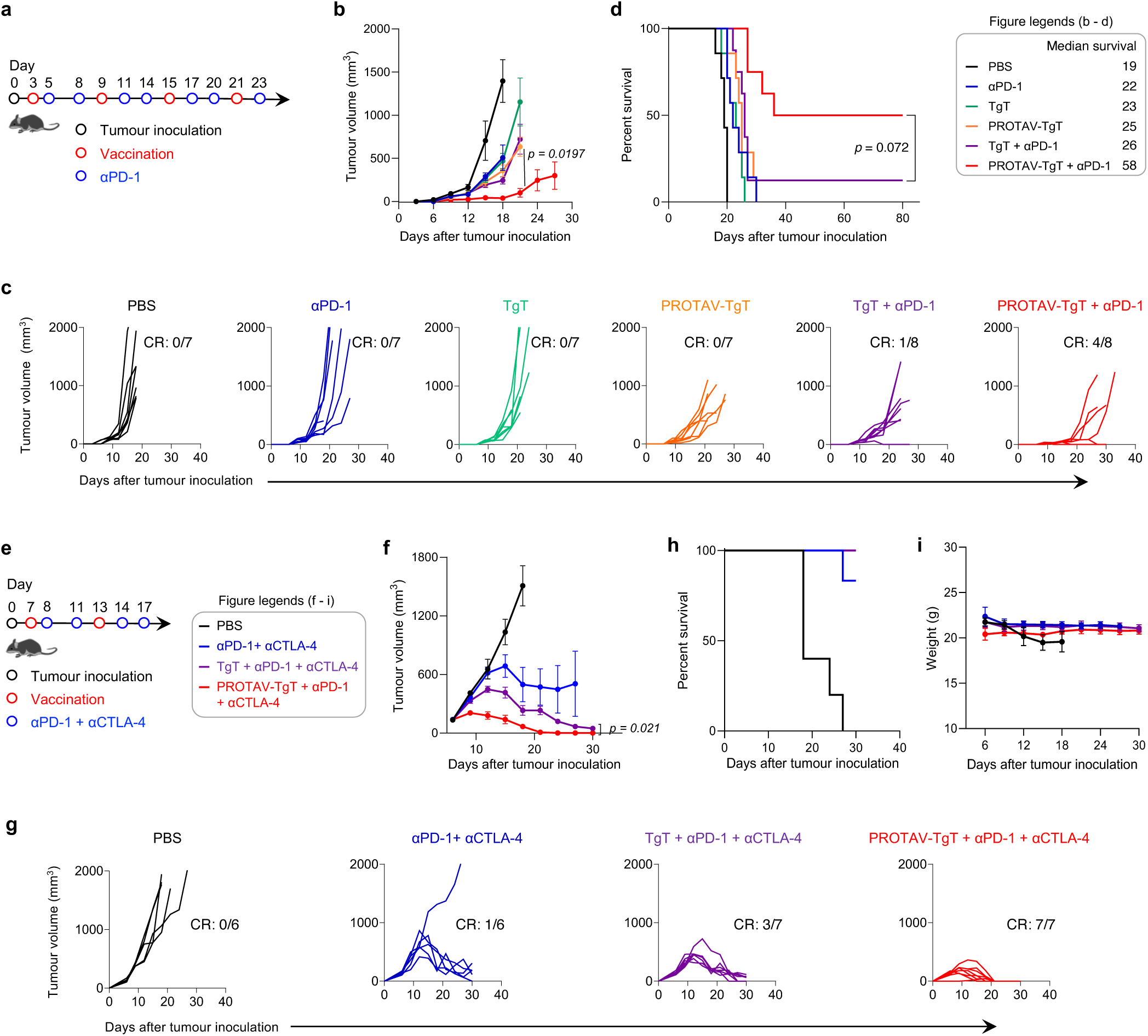
PROTAV-TgT, in combination with ICB, elicited systemic antitumor immunity for robust melanoma immunotherapy. **a**, Timeline for melanoma therapy study using PROTAV-TgT, alone or combined with αPD-1, in B16F10 melanoma-bearing C57BL/6 mice. B16F10 cells were *s.c*. inoculated into mice. Treatment started when the average tumor volumes reached 40 mm^3^. αPD-1: 150 μg, *i.p.* administration. **b-d**, Average B16F10 melanoma tumor growth curves (**b**), individual tumor growth curves (**c**), and Kaplan-Meier mouse survival curves of the above mice (**d**). Results showed that, relative to TgT, PROTAV-TgT, alone or combined with αPD-1, facilitated the tumor regression, promoted B16F10 melanoma therapeutic response, and improved mouse survival. **e**, Timeline for SM1 therapy study using PROTAV-TgT, alone or combined with αPD-1 and αCTLA-4, in *Braf^V600E^* SM1 melanoma-bearing C57BL/6 mice. SM1 tumors were first propagated in immunodeficient NSG mice, followed by tumor tissue excision and *s.c*. transplantation into syngeneic C57BL/6 mice. Treatment started when the average tumor volumes reached 150 mm^3^. αPD-1 and αCTLA-4: 100 μg each, *i.p.* administration. **f-g**, Average (**f**) and individual (**g**) SM1 tumor growth curves after the above treatments. **h-i**, Kaplan-Meier mouse survival (**h**) and mouse body weights (**i**) after the above treatments. Vaccines (PROTAV-TgT or TgT, Svg3, and CpG) were co-loaded in SM-102 LNPs and *s.c.* injected at mouse tail base (dose: 50 μg antigen, 2 nmole CpG, 1 nmole Svg3). CR: complete regression. Data represent mean ± s.e.m. (*n* = 6-8); statistical analysis was conducted using one-way ANOVA with Bonferroni post-test.

The robust melanoma therapeutic efficacy of PROTAV-TgT + ICB motivated us to test this regimen in a syngeneic mouse model of large SM1 *Braf^V600E^* melanoma. 50% human melanoma has *Braf^V600E^* mutation, which confers tumor cell resistance to apoptosis and poor prognosis.^60,61^ *Braf^V600E^* human melanoma can be closely recapitulated by murine *Braf^V600E^* melanoma models, such as SM1, for preclinical studies to faithfully predict clinical therapeutic efficacy.^61^ Moreover, relative to small tumors, large tumors are substantially more challenging for immunotherapy to eradicate, largely due to hostile TME and systemic immunosuppression, and the demand for immunotherapy to elicit a large quantity and good quality of antitumor immunity for tumor eradication. For optimal therapeutic outcome, dual ICIs of αPD-1 and cytotoxic T lymphocyte antigen-4 antibody (αCTLA-4) were used for combination immunotherapy with PROTAV-TgT vaccines. In C57BL/6 mice, seven days following tumor implantation (initial tumor volumes: 150 mm³), mice were treated with PROTAV-TgT, alone or combined with αPD-1 + αCTLA-4, with controls of PBS, αPD-1 + αCTLA-4, and TgT + αPD-1 + αCTLA-4 (**Fig. 6e**). As shown by tumor growth, PROTAV-TgT combined with αPD-1 + αCTLA-4 enhanced tumor growth inhibition relative to TgT + αPD-1 + αCTLA-4 and other control treatments (**Fig. 6f-g**). Notably, a single-dose PROTAV-TgT and ICB already showed significant tumor growth inhibition (day 9). This suggests that PROTAV-TgT elicits a rapid and potent antitumor immune response for combination with ICB. After only two doses of vaccines and four doses of ICB (day 21), PROTAV-TgT + αPD-1 + αCTLA-4 eradicated 100% of SM1 tumors; by contrast, at the same time point, none of the tumors was eradicated by any of the control treatments. While TgT + αPD-1 + αCTLA-4 showed gradual tumor growth inhibition and resulted in 3/7 tumor CR, the response was markedly less pronounced than that for PROTAV-TgT + αPD-1 + αCTLA-4. Lastly, during the treatment course, PROTAV-TgT + αPD-1 + αCTLA-4 did not cause any significant mouse body weight changes in SM1 (**Fig. 6i**). Altogether, these findings underscore the ability of PROTAV to potentiate antitumor T cell responses for robust tumor immunotherapy in combination with ICB.

## Discussion

Cancer therapeutic vaccines can elicit or expand antitumor CD8^+^ T cells that are pivotal for the combination immunotherapy of various types of cancers, such as melanoma. The antigens in cancer therapeutic vaccines often need to be proteolytically processed for MHC binding, antigen presentation, thereby priming naïve T cells or expanding pre-existing antigen-specific T cells. We developed PROTAVs as a novel platform to facilitate proteolysis and thus promote antitumor T cell responses for cancer immunotherapy. Mechanistically, the ability of vaccines to elicit antitumor T cell responses was promoted by ligand-medicated E3 ligase engagement that promotes antigen ubiquitination, proteolytic processing into minimal antigenic epitopes, and cross-presentation of the resulting antigenic epitopes to CD8^+^ T cells. PROTAV promotes the proteolysis of exogenous protein/peptide antigens, in comparison to PROTAC as an emerging platform for the degradation of endogenous intracellular proteins. We expect that the research and development of PROTAC over the past two decades would facilitate the research and development of PROTAVs. PROTAVs hold the potential for rapid development and clinical translation, as supported by the potential of cancer therapeutic vaccines, the vast unmet clinical need for novel and effective cancer immunotherapeutic approaches, and the clinically proven efficacy and safety of PROTACs for molecular ligand-promoted proteolysis in multiple clinical trials. Advanced melanoma is a type of aggressive tumor which has benefited from recent advancement of immunotherapy, including ICB and adoptive cell transfer therapies. However, current immunotherapies for melanoma still have the limitations, such as limited durable response rates and low affordability. Cancer therapeutic vaccines, including peptide vaccines targeting multivalent tumor-associated antigens and neoantigens, hold the potential to be combined with current immunotherapies for optimal therapeutic outcomes to maximize their therapeutic benefits. PROTAVs can be easily adopted by current peptide or protein vaccines and further promote anti-tumor T cell responses against these antigens.

With a modular structure that can be easily synthesized via biofriendly conjugation, PROTAVs allow for easy adjustment of E3 ligands, linkers, and various protein/peptide antigens. In this study, we used CRBN-bindign pomalidomide as a representative E3 ligand. Pomalidomide is an FDA-approved drug interacting with CRBN and inducing degradation of essential Ikaros transcription factors. Physically mixing pomalidomide with OVA did not significantly impact the anti-OVA T cell responses, ruling out the confounding intrinsic immunomodulatory effect of pomalidomide in PROTAV. We expect that the clearly understood pharmacology, toxicology, and safety profile of pomalidomide, as well as ongoing clinical studies of pomalidomide-based PROTACs, would facilitate the clinical translation of PROTAV. Moreover, alternative E3 ligands (e.g., VH032 for Von Hippel-Lindau (VHL), cIAP1 for inhibitor of apoptosis proteins (IAP))^33^ can be readily tested in PROTAV. Site-specific conjugation of E3 ligands on antigens can be desirable to not only avoid conjugating these ligands on antigenic minimal epitopes and affecting the MHC binding ability of these epitopes, but also for reproducible batch-to-batch manufacturing. Here, we explored pomalidomide conjugation on natural amino acids, primarily lysine and cysteine. While lysine conjugation may lead to competition for the use of lysine between E3 ligand conjugation and ubiquitination, maleimide-cysteine conjugation may bypass such complication. Moreover, we envision that unnatural amino acids can be site-specifically incorporated into protein/peptide antigens, allowing for bioorthogonal site-specific conjugation of E3 ligand on unnatural amino acids in antigens without affecting any minimal antigenic epitopes^62,63^.

PROTAVs could have broad applications in proteins or synthetic long peptide vaccines. Relatively to large proteins, synthetic long peptides contain minimal peptide epitopes to elicit antitumor T cell responses and can have the advantage of quick large-scale manufacturing and high pharmaceutical stability. Relative to minimal peptide epitopes, synthetic long peptide antigens allow de novo antigen processing in APCs for potent and durable immunomodulation with minimal tolerance, the latter of which can be caused by MHC presentation of antigen epitopes from APCs to T cells in the absence of co-stimulation or proinflammatory cytokines.^40,41^ Further, the simple and efficient biocompatible conjugation of E3 ligands on tumor antigens permits their easy adoption into current protein/peptide vaccines, such as tumor-associated antigens, tumor neoantigens, and oncoviral antigens. Multivalent antigen vaccines are poised to be used for tumor immunotherapy because 1) they could overcome the complications associated with the vast intra-patient and inter-patient tumor antigenic heterogeneity and the resulting tumor immune escape, and 2) the use of both MHC-I and MHC-II classes of antigens yields the optimal tumor therapeutic efficacy. To leverage the benefits of multivalent antigen vaccines, multivalent antigens need proteolytic processing to generate minimal antigen epitopes to bind MHC and activate T cell responses.

The overall weak immunogenicity of tumor antigens and the systemic immunosuppression in cancer patients demands the use of potent immunostimulant adjuvants for vaccines to elicit robust systemic antitumor immunity while reducing the systemic and intratumoral immunosuppression. Previously, we engineered Svg3 oligonucleotide as a potent and selective cGAS agonist that can elicit type-I IFN responses, which are instrumental to eliciting antitumor T cell responses;^25^ using two TLR agonists, we further showed that biadjuvant can promote T cell responses for tumor immunotherapy.^64^ Here, we combined Svg3 and CpG as bi-adjuvant to further promote the ability of PROTAV to elicit antitumor immunity. The cationic charges of Svg3 and CpG oligonucleotides as well as the high hydrophobicity of many antigenic peptides, especially when conjugated with pomalidomide, allows for easy co-loading into LNPs. Although ionizable LNPs are immunostimulatory to elicit potent antibody responses,^65^ their ability to potentiate T cell responses can be further improved. Our study also suggests the potential of CpG and Svg3 as combination adjuvants to potentiate LNP-based vaccines to elicit T cell responses. Taken together, these results support the potential of PROTAVs as a novel platform of cancer therapeutic vaccines for broad application in the robust immunotherapy of many types of cancer.

## Methods

### PROTAV synthesis and characterization

*Pomalidomide-linker-NHS*. A series of linkers (e.g., EG_4_) was used in pomalidomide-linker-COOH (Tocris Bioscience, Minneapolis, MN) as starting materials. Pomalidomide-linker-COOH (2.0 mg, 3.8 μmol), NHS (0.5 mg, 4.3 μmol), and EDC hydrochloride (3.9 mg, 20.3 μmol) were added into a glass vial with 100 μL anhydrous dimethylformamide (DMF) and stirred at room temperature for 2 hours. After that, DMF was evaporated and 500 μL chloroform was added into the vial. The mixture was washed with H_2_O twice, the organic phase was collected, and the solvent was removed under vacuum to obtain pomalidomide-linker-NHS. Then pomalidomide-linker-NHS (0.8 mg), antigen peptide or protein (11.0 mg) were added into a glass vial with 1 mL phosphate-buffered saline (pH 8.0) and stirred at room temperature for 2 hours. After that, the PROTAV was purified by ultrafiltration (10-kDa cutoff; Millipore). Fluorescence spectra (BioTek Cytation 5; with fluorescamine as a probe)^66^ or LC-MS (Thermo Scientific) was used to analyze the copy numbers of pomalidomide-linker conjugated per antigen peptide or protein molecule.

*Pomalidomide-EG_4_-maleimide*. Pomalidomide-EG_4_-NH_2_ (2 mg, 3.7 μmol) and maleic anhydride (0.7 mg, 7.1 μmol) were added into a glass flask with 1 mL chloroform and refluxed with stirring (60 °C, 24 h). Then, chloroform was evaporated, and acetic anhydride (1 mL) and sodium acetate (0.7 mg, 7.5 μmol) were added into the flask. The reaction solution was refluxed with stirring (90 °C, 24 h). Next, the reaction solution was washed with 1 M aqueous sodium hydroxide solution. The organic phase was extracted by dichloromethane. The solvent was removed under vacuum to obtain pomalidomide-EG_4_-maleimide. Next, pomalidomide-EG_4_-Mal (0.8 mg) and peptide/protein antigen (11.0 mg) were added into a glass vial with 1 mL H_2_O (pH 7.0) and stirred at room temperature (4 h). Lastly, PROTAV was purified by ultrafiltration (10 kDa cutoff; Millipore) and verified by UV-vis and LC-MS.

### Nano ITC measurement of PROTAV-CRBN binding

The binding affinity assay was performed using NanoITC (TA Instruments). Incremental titration (20, 2.5 μL injections in every 300 s) of CRBN protein (7.5 μΜ) into PROTAV-OVA (100 μΜ) was used for the assay. Background titration was performed using same titration method to subtract the heat from protein buffer. The ITC thermogram was analyzed using NanoAnalyze, and the Nano ITC data was fit into a one site-specific binding model in GraphPad to determine the *K_d_*.

### Plasma stability study of pomalidomide-EG_4_-COOH

The *in vitro* stability of pomalidomide-EG_4_-COOH was studied in CD1 mouse plasma (Innovative Research Inc, Novi, US). Pomalidomide-EG_4_-COOH powder was weighed and dissolved in DMSO to a working concentration of 100 μM. 495 μL of pre-warmed plasma at 37 °C was prepared followed by spiking 5 μL of the 100 μM pomalidomide-EG_4_-COOH working solution, to a final concentration of 1 μM. The spiked mouse plasma was incubated at 37 °C for 4 h. 40 μL plasma samples were separately acquired at 0, 5, 10, 15, 30, 45, 60, 120, and 240 min, and reactions were rapidly terminated by adding 160 μL of cold acetonitrile containing internal standard (10 ng/mL IPI549 in acetonitrile) at each time point. After quenching, the samples were shaken for 5 min at 700 rpm and centrifuged at 4000 rpm for 10 min. 100 μL of the supernatant from each well was transferred into a 96-well sample plate for LC/MS analysis. The *in vitro* plasma half-life (t_1/2_) was calculated using the expression t_1/2_ = 0.693/b, where b is the slope found in the linear fit of the natural logarithm of the fraction remaining of the test compound vs. incubation time.

### Western blotting (WB)

WB was employed to investigate the intracellular degradation of PROTAV-OVA. DC2.4 cells were treated with PROTAV-OVA or OVA. Subsequently, the cells were harvested and lysed in RIPA lysis buffer (comprising 50 mM Tris-HCl, pH 7.4, 150 mM NaCl, 1% NP-40, and 0.1% SDS), with the addition of 1 mM phenylmethylsulfonyl fluoride to inhibit protease activity. The lysates were then centrifuged at 12,000 g for 10 minutes at 4°C, and the supernatants were collected as the protein fraction. Protein concentrations were measured using a BCA protein assay kit, after which the samples were denatured at 70 °C for 15 minutes in the presence of 25% (v/v) loading buffer. Proteins were subsequently separated *via* SDS-polyacrylamide gel electrophoresis (SDS-PAGE) and transferred to a polyvinylidene fluoride (PVDF) membrane (Merck Millipore). To prevent non-specific binding, the membrane was blocked with 5% non-fat dry milk in Tris-Buffered Saline with Tween 20 (TBST) for 1 hour at room temperature, followed by overnight incubation at 4°C with the primary antibody, diluted 1:1000 in 5% non-fat dry milk. The samples were then incubated with an HRP-conjugated secondary antibody, diluted 1:5000 in 5% non-fat dry milk, for 1 hour at room temperature. OVA was analyzed using a BioRad GelDoc imaging system.

### Co-IP

Co-IP was used to evaluate the interaction between OVA and ubiquitin. DC2.4 cells were treated with PROTAV-OVA and OVA, respectively, for 12 - 16 h. Proteins were extracted from as-treated cells using a magnetic IP/co-IP kit (Thermo Fisher Scientific, Waltham, MA). The protein lysates were incubated with anti-OVA magnetic beads for 2 - 3 h. The beads were then washed for four times using a buffer of 50 mM Tris-HCl (pH 7.5), 150 mM NaCl, 20% glycerol, 0.1% Triton X-100, and 1 mM EDTA (pH 8.0). The immunoprecipitates were resolved via SDS-PAGE, transferred to a PVDF membrane (Merck Millipore), and subsequently probed with anti-ubiquitin antibodies. Ubiquitin analysis on the resulting gel was detected using a BioRad GelDoc imaging system.

### PROTAV LNP preparation

PROTAV LNPs were synthesized using microfluidic chips on a NanoGenerator Flex-S instrument (PreciGenome, San Jose, CA). An ethanol phase containing peptides and SM-102 lipid, DSPE, DMG-PEG_2000_, and cholesterol (the lipid molar ratio is 50: 10: 1.5: 38.5) was mixed with an aqueous phase of CpG or Svg3 in 10 mM citrate buffer at a flow rate of 2 mL/min. PROTAV LNPs were dialyzed in 1x PBS at 4 °C for 12 hours using a microdialysis cassette (20,000 MWCO, Thermo Fisher Scientific, Waltham, MA). The diameters and polydispersity index (PDI) of the LNPs were measured with a Zetasizer Nano (Malvern Instruments, Malvern, U.K.).

### Cryo-EM of LNPs

LNPs were subject to dialysis and concentration to ∼90 mg/mL of lipid using Amicon ultracentrifugation filters. Grids for cryo-EM imaging of TgT-LNPs and blank LNPs were prepared by applying 6 μL purified encapsuling sample at a concentration of 10 mg/ml to a glow-discharged (1 min at 5 mA) Ultrathin Carbon on Quantifoil (R1.2/1.3, 300 mesh, Cu) (Ted Pella, Redding, CA). This was followed by plunge-freezing in liquid ethane using an FEI Vitribot (100% humidity, temperature: 4°C) in two steps: 1) with blot force of 0, wait time of 10 s, and 2) with blot force of 0, wait time of 0 s. Grids were clipped and stored in liquid nitrogen until data collection. Cryo-EM images were collected at the University of Michigan Cryo-EM Facility on a Talos Arctica transmission electron microscope (Thermo Fisher Scientific, Waltham, MA) with K2 Summit direct detector (Gatan, Pleasanton, CA) operating at 200 kV.

### Cell culture

All cells were cultured in medium (Gibco, Brooklyn, NY) supplemented with 10% heat-inactivated FBS, 100 U/mL penicillin, and 100 μg/mL streptomycin, at 37 °C with 5% CO_2_. EG7.OVA cells were cultured in RPMI-1640 medium additionally supplemented with 2 mM L-glutamine and 0.4 mg/mL G418. BMDCs, OT-1 cells, and DC2.4 cells were cultured in RPMI-1640 medium additionally supplemented with 2 mM l-glutamine, 50 μM 2-mercaptoethanol, 1× non-essential amino acids and 10 mM HEPES. B16F10 cells were cultured in DMEM medium.

### BMDC isolation and culture

BMDCs were obtained from 6–8 week old C57BL/6 mice. Femurs and tibias were harvested and cut at the epiphyses, and their bone marrow was flushed out three times with 10 mL RPMI-1640 medium (Gibco, Brooklyn, NY). Cell suspensions were filtered through 70-μm cell strainers (BD Falcon, Franklin Lakes, NJ), centrifuged, and treated with ACK buffer (Gibco, Brooklyn, NY) for lysis. Cell pellets were resuspended in RPMI-1640 medium (10% FBS and 20 ng/mL GM-CSF) to achieve a density of 2 × 10^6^ viable cells per 75-mm petri dish. On day 6, non-adherent and loosely adherent cells were collected as BMDCs by gently washing with PBS and pooled for further studies.

### Flow cytometry

Immunostained cultured cells, mouse blood cells, and single-cell suspensions from tumor tissue and lymph node homogenates were analyzed by flow cytometry (Cytek Aurora, Cytek Biosciences, Fremont, CA). The antibodies used for immunostaining are detailed in **Supplementary Table 3**. Data were analyzed using FlowJo V10 software, and representative gating strategies for flow cytometry data analysis are shown in **Supplementary Fig. 14-17**.

### *In vitro* and *in vivo* antigen SIINFEKL presentation on DCs

For in vitro cell study, 2 × 10^5^ DCs were seeded in 24-well plates and treated with PROTAV-OVA, OVA, and controls, respectively, for 24 hours. Harvested DCs were then stained with an APC-labeled anti-mouse H-2K^b^/SIINFEKL complex antibody (Biolegend). For in vivo studies, C57BL/6 mice were *s.c.* administered with PROTAV-OVA or OVA (10 μg), and PBS, respectively, with CpG (2 nmole) (Integrated DNA Technologies, Coralville, IW) as adjuvant. After a series of time, draining inguinal lymph nodes were harvested and homogenated to prepare single cells. H-2K^b^/SIINFEKL complex on DCs was stained as above in the resulting single cells. Cells were analyzed using flow cytometry.

### OT-1 CD8^+^ T cell activation

OT-1 cells were isolated from OT-1 mice (Jackson Laboratory) using CD8^+^ T cell isolation kit (Miltenyi, Auburn, CA). OT-1 cells were then stained with an intracellular dye CFSE, followed by washing to remove free CFSE dyes. DCs were treated with PROTAV-OVA, OVA, or controls. Next, CFSE-stained OT-1 cells and as-treated DCs with a ratio of 20:1 were co-cultured for up to three days. Upon recognition of SIINFEKL antigen presented on DCs, SIINFEKL-specific OT-1 cells would be activated and proliferate. The OT-1 cell activation can be measured via the CFSE fluorescence signal dilution by flow cytometry.

### Intracellular delivery of PROTAV

*In vitro* intracellular delivery of PROTAV was studied using dye-labeled PROTAV-OVA by confocal microscopy and flow cytometry. PROTAV-OVA was labeled with FITC using NHS-FITC and PROTAV-OVA that was synthesized by maleimide-thiol conjugation of OVA and pomalidomide-EG_4_. FITC-PROTAV-OVA and FITC-OVA, respectively, were incubated with DC2.4 cells for up to 24 h, prior to staining with Lysotracker Green DND-26 (Life Technologies, Carlsbad, CA) and Hoechst 33342 (Life Technologies, Carlsbad, CA) according to manufacturers’ instructions. Cells were then washed before confocal microscopy (LSM 780, Zeiss, Chesterfield, VA). Cells treated as above were also analyzed by flow cytometric analysis to measure the FITC fluorescence intensity of cells.

### RNA-seq

Total RNA extraction was achieved using the AllPrep DNA/RNA Mini Kit (QIAGEN) and included in-column DNase I digestion to ensure the RNA was free of DNA contamination. The extracted RNA’s integrity was checked with the Agilent RNA 6000 Pico Kit (Agilent). For mRNA-seq library preparation, 1 μg of total RNA (with an RIN score above 8) was used, according to Illumina stranded mRNA-seq preparation Guide. In brief, total RNA was processed to isolate mRNA using oligo (dT) magnetic beads, which then underwent cation fragmentation and reverse transcription into complementary DNA (cDNA). The cDNA was end-repaired, ligated to RNA index anchors, and PCR-amplified with random primers to generate indexed libraries, using IDT for Illumina UD indexes. After ensuring the quality, these barcoded libraries were equally pooled and paired-end sequenced (2 x 150 bp) on the NextSeq 2000 instrument (Illumina). FastQC v0.11.9 was used to evaluate the quality of RNA-Seq reads. The STAR aligner version 2.7.6a^67^ was employed to align reads from individual samples to the mouse reference genome GRCm39. FeatureCounts^68^ was used to aggregate raw gene counts of the aligned reads. Differential gene expression analysis was conducted using the Bioconductor package DESeq2 v1.30.0^69^ with normalized and filtered gene counts from the RNA-Seq data. Genes exhibiting differential expression with an adjusted *p* value of less than 0.05 were further analyzed for Gene Ontology and KEGG pathway enrichment analyses using the ORA.

### Animal studies

All animal studies were conducted in compliance with the Guide for the Care and Use of Animals under protocols approved by the Institutional Animal Care & Use Committee (IACUC) at University of Michigan and Virginia Commonwealth University.

### PBMC T cell responses

*Tetramer staining, immune memory, and immune checkpoint expression levels on T cells.* Peripheral blood was drawn from vaccinated mice. ACK lysis buffer was used to lyse red blood cells for 10 minutes at room temperature. Any blood clots were filtered out, and the cells were washed twice with PBS. Zombie Aqua™ Fixable Viability Kit (BioLegend, San Diego, CA) was used for live-dead cell staining for 20 minutes at room temperature, followed by cell washing with FCS buffer (PBS with 0.1% FBS). Cells were blocked with αCD16/CD32 (BioLegend, San Diego, CA) for 10 min, and were then incubated with a staining cocktail containing anti-mouse CD3-PerCP-Cy5.5, anti-mouse CD8α-APC-Cy7, CD44-AF647, CD62L-FITC, anti-mouse PD-1-BV421, anti-mouse Tim-3-BV605, and PE-conjugated tetramers (NIH Tetramer Core Facility). After 30 minutes at room temperature, cells were washed, and then fixed using 100 μL Cytofix per well of 96-well plates at 4°C for 20 minutes. Finally, cells were washed and resuspended with Perm/Wash buffer for analysis by flow cytometry. For immune memory analysis, cells were gated based on CD44^high^CD62L^high^ central memory CD8^+^ T cells, CD44^high^CD62L^low^ effector memory CD8^+^ cells, and CD44^low-med^CD62L^high^ naive CD8^+^ T cells.

*Intracellular cytokine staining of T cells.* Intracellular cytokine staining in T cells was performed as before.^9^ Briefly, peripheral blood from immunized mice was collected and processed as above to obtain lymphocytes and lyse red blood cells. The lymphocytes were then transferred into U-bottom 96-well plates containing 200 μL of T cell culture medium. Cells were restimulated by 40 μg/mL antigen peptides for 4 hours (CS Bio, Menlo Park, CA; peptide sequences in **Supplementary Table 2**), after which GolgiPlug Protein Transport Inhibitor with brefeldin A (Thermo Fisher Scientific) was added. The cells were then incubated for 6 hours prior to αCD16/CD32 staining at room temperature for 10 minutes. Following this, cells were stained with anti-mouse CD8α-APC-Cy7, anti-mouse CD4-PerCP-Cy5.5, and the Zombie Aqua™ Fixable Viability Kit (BioLegend, San Diego, CA) (for 20 minutes at room temperature). Cells were then washed and fixed using Cytofix (BD Biosciences), and were permeabilized with 200 μL Cytoperm solution (BD Biosciences), followed by washing with Perm/Wash Buffer (BD Biosciences). Upon permeabilization, cells were stained with PE anti-mouse IFN-γ and FITC anti-mouse TNF-α (BioLegend). After final washing, the cells were prepared for flow cytometric analysis.

### Immune analysis of TME, spleen, and lymph nodes in tumor-bearing mice

Tumors were established by *s.c*. injection of tumor cells at mouse shoulders. Tumor-bearing mice were treated with PROTAV or control vaccines, alone or in combination with ICIs. Blood and tissues of interest were collected four days after the last treatment. Tissues were weighed and dissected into small pieces prior to digestion treatment in RPMI 1640 media (15 min, 37 °C) supplemented with collagenase (Sigma, St Louis, MO) and DNase I (New England Biolabs, Ipswich, MA). Tissues were filtered through a 70 μm cell strainer (Fisher Scientific, Pittsburg, PA) and treated with ACK Lysing Buffer (Gibco, Brooklyn, NY) for red blood cell lysis. The resulting single cells were then washed and stained with live-dead staining dye, followed by staining with antibody cocktails and tetramers according to the manufacturer’s instructions. At the end of staining, cells were washed and fixed prior to flow cytometric analysis as described above.

### Tumor immunotherapy

Female C57BL/6 mice (6–8 weeks old, Charles River Laboratories, *n* = 6–10) were *s.c.* injected on the shoulder with 3 × 10^5^ tumor cells (i.e., B16F10, SM1). Tumor progression was assessed using caliper measurements. Mice were treated when tumor reached a certain volume (e.g., 50 mm^3^ B16F10 tumors or 150 mm^3^ SM1 tumors). SM1 tumor cells were thawed and directly injected into NSG mice for tumor expansion. After 15 days, tumors were excised at an average volume of approximately 1000 mm^3^. The harvested tumors were dissected into small cubes (1-2 mm^3^), mixed with Matrigel (Corning, USA) at a 1:1 ratio, and were subcutaneously implanted into the right hind limbs of C57BL/6 mice. Vaccines were administered subcutaneously at the tail base for optimal lymphatic drainage (10-50 μg antigen, 2 nmole CpG and/or 1 nmole Svg3, and 100 μg αPD-1 or αCTLA-4 (Bio X Cell, Inc., NH). αPD-1 or αCTLA-4 was *i.p.* administered. Tumor dimensions and mouse weights were recorded every 3 days. Mice were euthanized if the largest tumor dimension reached 2 cm, tumor volume surpassed 2000 mm^3^, or ulceration occurred. Tumor volumes were calculated as (length x width^2^)/2 and analyzed as previously described, and results were processed using GraphPad Prism 7 (La Jolla, CA).

## Statistical analysis

Data represent mean ± s.e.m., unless denoted otherwise. Statistical analysis was performed in GraphPad Prism Software version 5.0. *P*-values were calculated by one-way ANOVA with Bonferroni post-test, unless denoted otherwise.

## Reporting Summary

Further information on research design is available in the Nature Research Reporting Summary linked to this article.

## Data availability

The data supporting the results in this study are available within the paper and its Supplementary Information. Source data for the figures will be provided with this paper.

## Acknowledgements

G.Z. acknowledges funding support from NIH (R01CA286122, R01CA266981, R01AI168684, R35GM143014), DoD CDMRP Breast Cancer Breakthrough Award Level II (BC210931/P1), American Cancer Society Research Scholar Grant (RSG-22-055-01-IBCD), University of Michigan Rogel Cancer Center Discovery Award. F.C. acknowledges fund support by the National Natural Science Foundation of China (82102203) and Shanghai Pujiang Program (23PJ1411300). Research reported in this publication was supported by the National Cancer Institute under Award Number P30 CA046592 by the use of the following Cancer Center Shared Resource: Flow Cytometry, Immune Monitoring, Pharmacokinetics, and Proteomics. We thank the University of Michigan ULAM (Unit for Laboratory Animal Medicine) In Vivo Animal Core (IVAC) and the Center for Structural Biology for providing service. We thank the Genomics Core, and Bioinformatics Shared Resource and Flow Cytometry Shared Resource at Massey Comprehensive Cancer Center (MCCC) at Virginia Commonwealth University for providing service. MCCC is supported, in part, with funding from NIH-NCI Cancer Center Support Grant P30 CA016059. We thank NIH Tetramer Core for kindly providing tetramer reagents. The content is solely the responsibility of the authors and does not necessarily represent the official views of the National Institutes of Health.

## Author contributions

G.Z. conceptualized the project. G.Z., T.S., and F.C. initiated the project. G.Z., T.S., F.C., and Q.W. designed the project. T.S., F.C., Q.W., X.L., S.Z., M.W., Y.X., S.L., J.D., G.X., C.Y., H.W., W.Z., B.W., performed experiments. R.T., K.M.T., J.Z. analyzed RNAseq data. G.Z., Q.W., T.S., F.C., D.X., and S.W. analyzed data. G.Z. and Q.W. drafted the manuscript. All authors reviewed the manuscript.

## Supplementary Information

**Supplementary Fig. 1.**
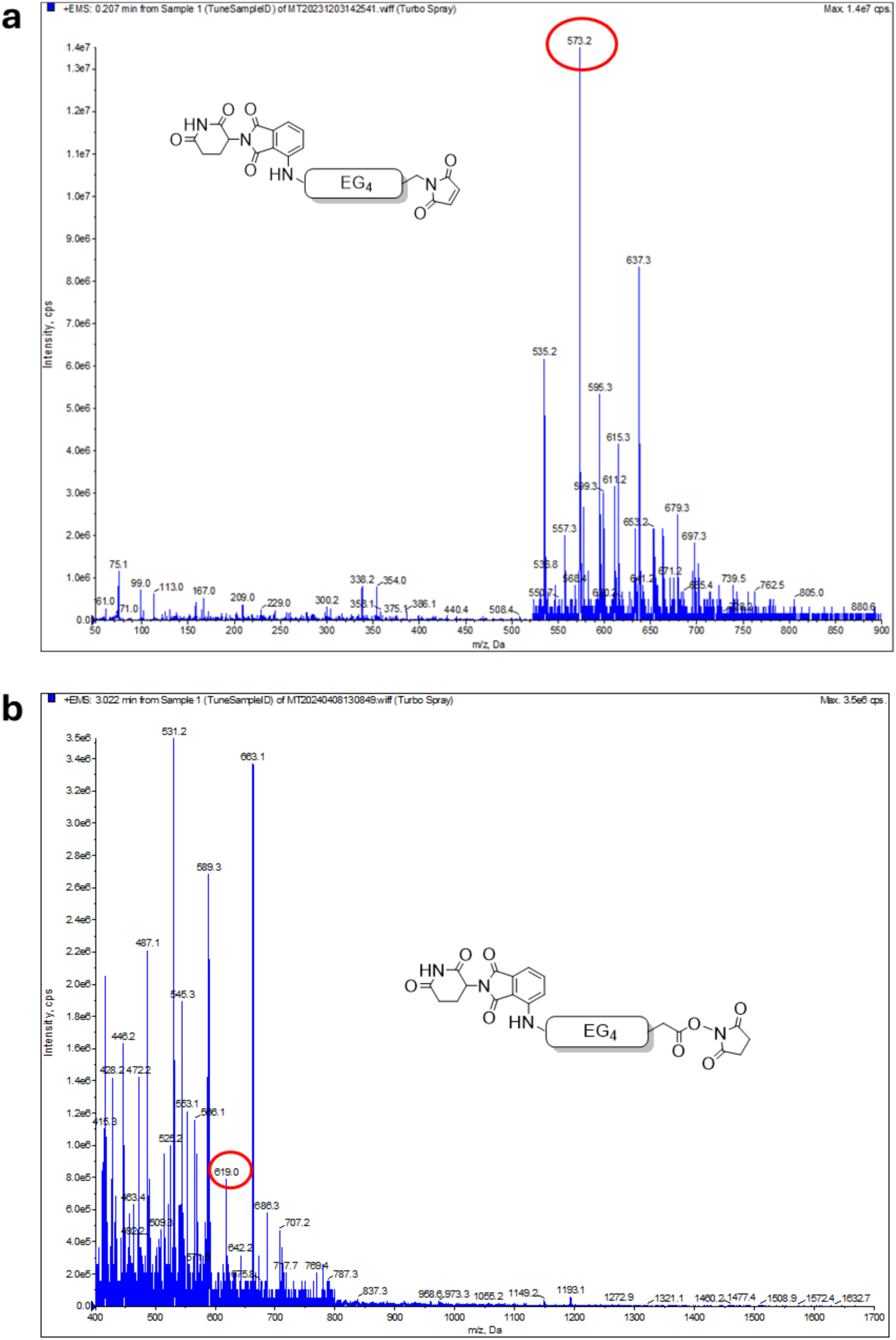
Chemical structures and ESI MS spectra of pomalidomide-EG_4_-Mal (a) and pomalidomide-EG_4_-NHS (b).

**Supplementary Fig. 2.**
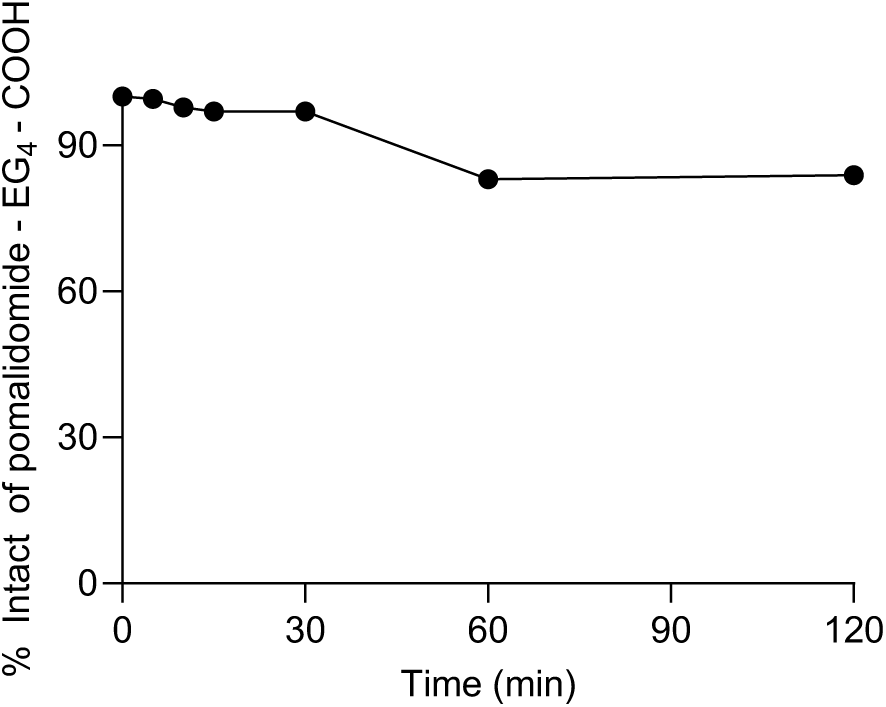
Stability of pomalidomide-EG_4_-COOH in serum measured by LC-MS. Pomalidomide-EG_4_-COOH was incubated in serum for a series of durations up to 2 h, followed by LC-MS measurement to quantify the remaining intact pomalidomide-EG_4_-COOH.

**Supplementary Fig. 3.**
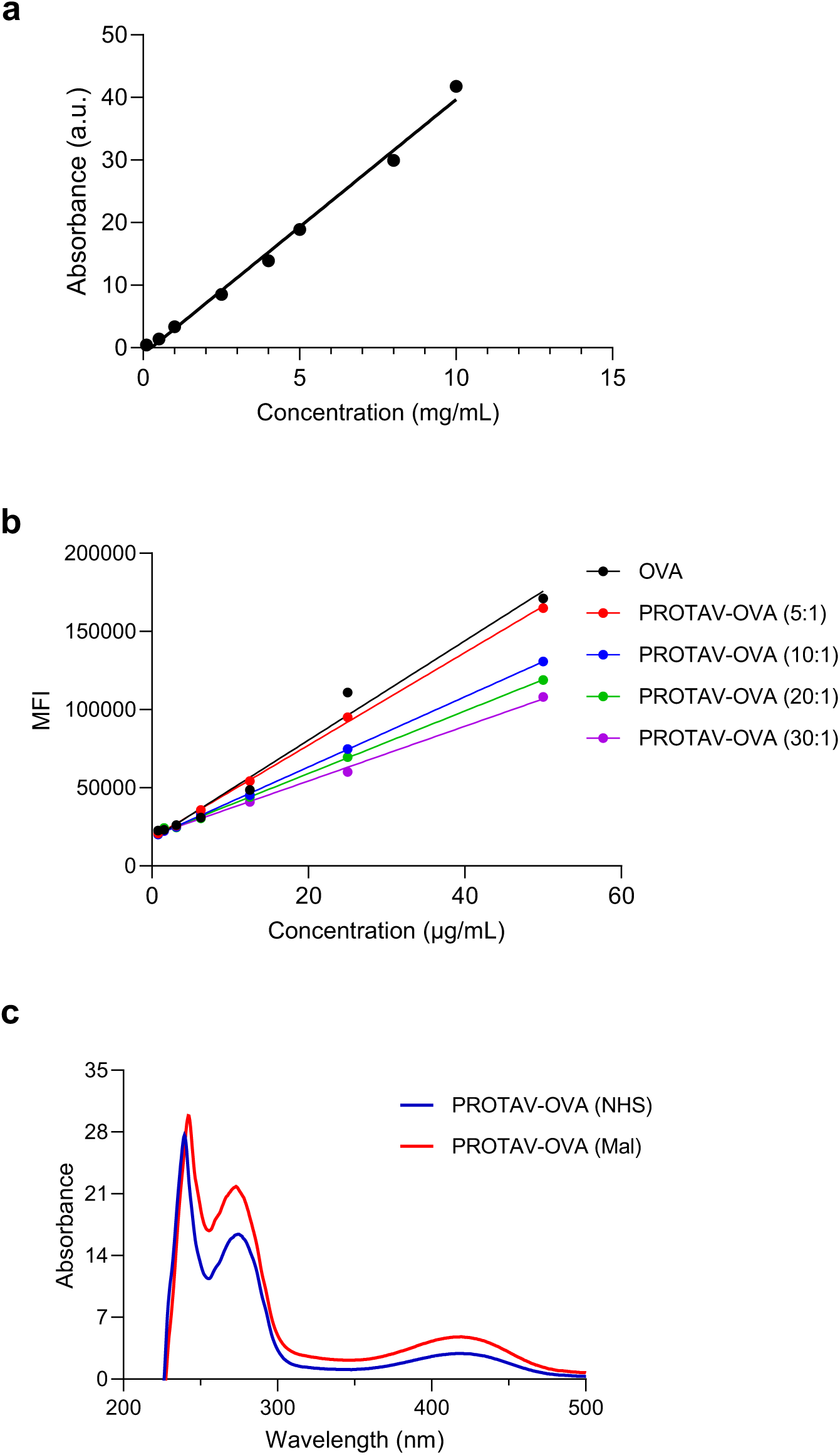
Characterization of PTOTAV-OVA conjugation ratio. **a**, Standard curve of pomalidomide-EG_4_ absorbance (420 nm) *vs*. concentration. **b**, MFI of amine-reactive beads used in an amine-reactive fluorescent assay. The results were used to determine the copy numbers of pomalidomide-EG_4_-NHS conjugated per OVA with a series of pomalidomide-EG_4_-NHS: OVA feeding molar ratios. **c**, UV-vis spectra of PROVA-OVA with two synthesis methods *via* NHS-EDC and maleimide-thiol conjugations, respectively.

**Supplementary Fig. 4.**
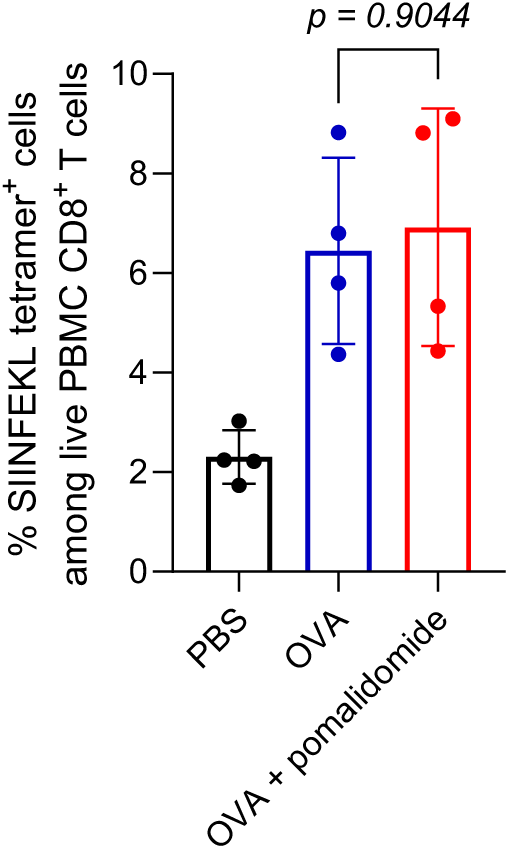
Tetramer staining of mouse PBMC CD8^+^ T cells from mice immunized with OVA or physical mixture of OVA and pomalidomide (day 21). C57BL/6 mice (6-8 weeks; *n* = 4) were immunized by *s.c*. administration at tail base (day 0, day 14). CpG (2 nmole) adjuvant was mixed with OVA or pomalidomide + OVA (10 μg).

**Supplementary Fig. 5.**
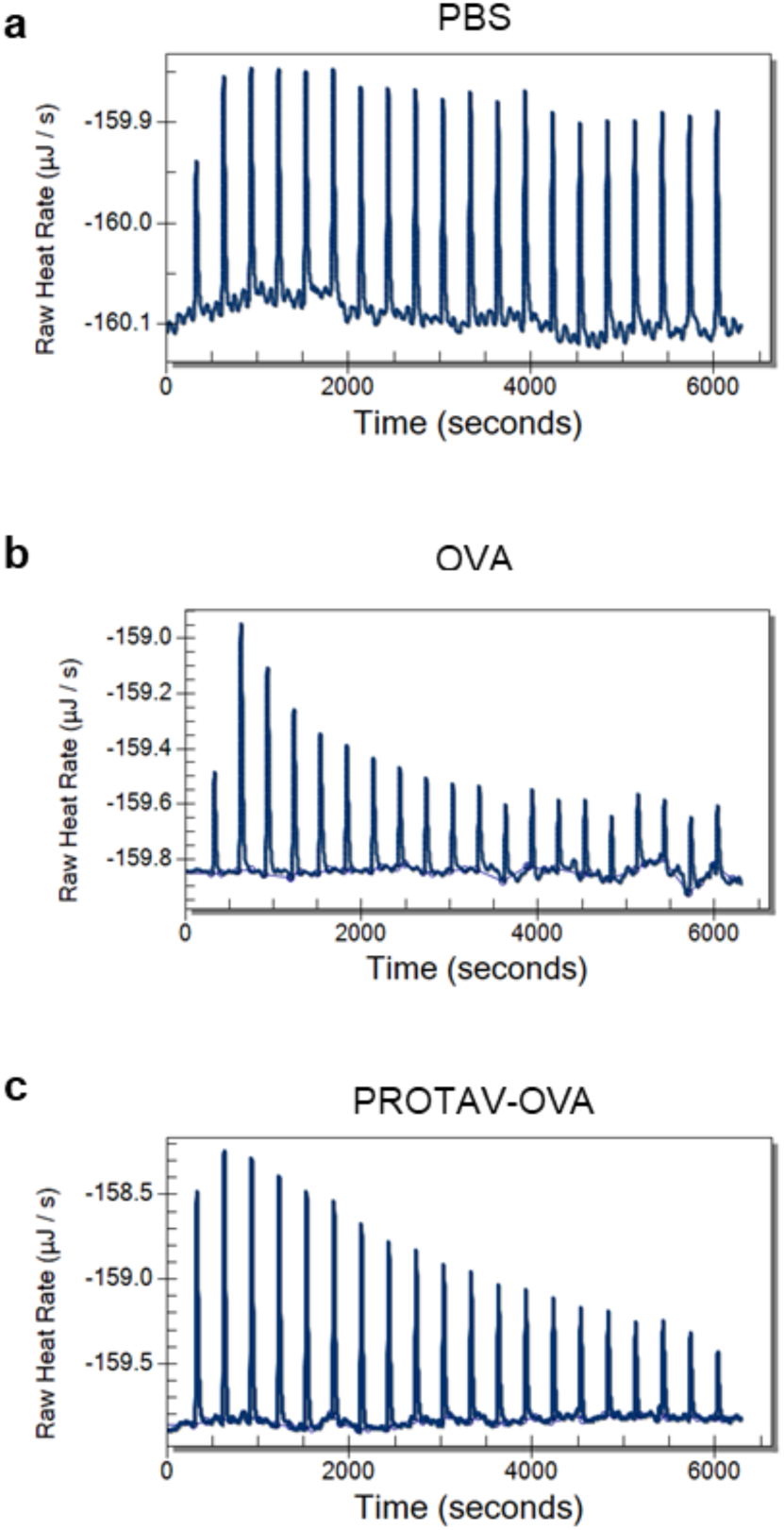
Nano ITC results showing the CRBN binding between PROTAV-OVA and CRBN protein. Shown are nano ITC graphs of binding kinetics between CRBN and PBS (**a**), OVA (**b**), and PROTAV- OVA (**c**).

**Supplementary Fig. 6.**
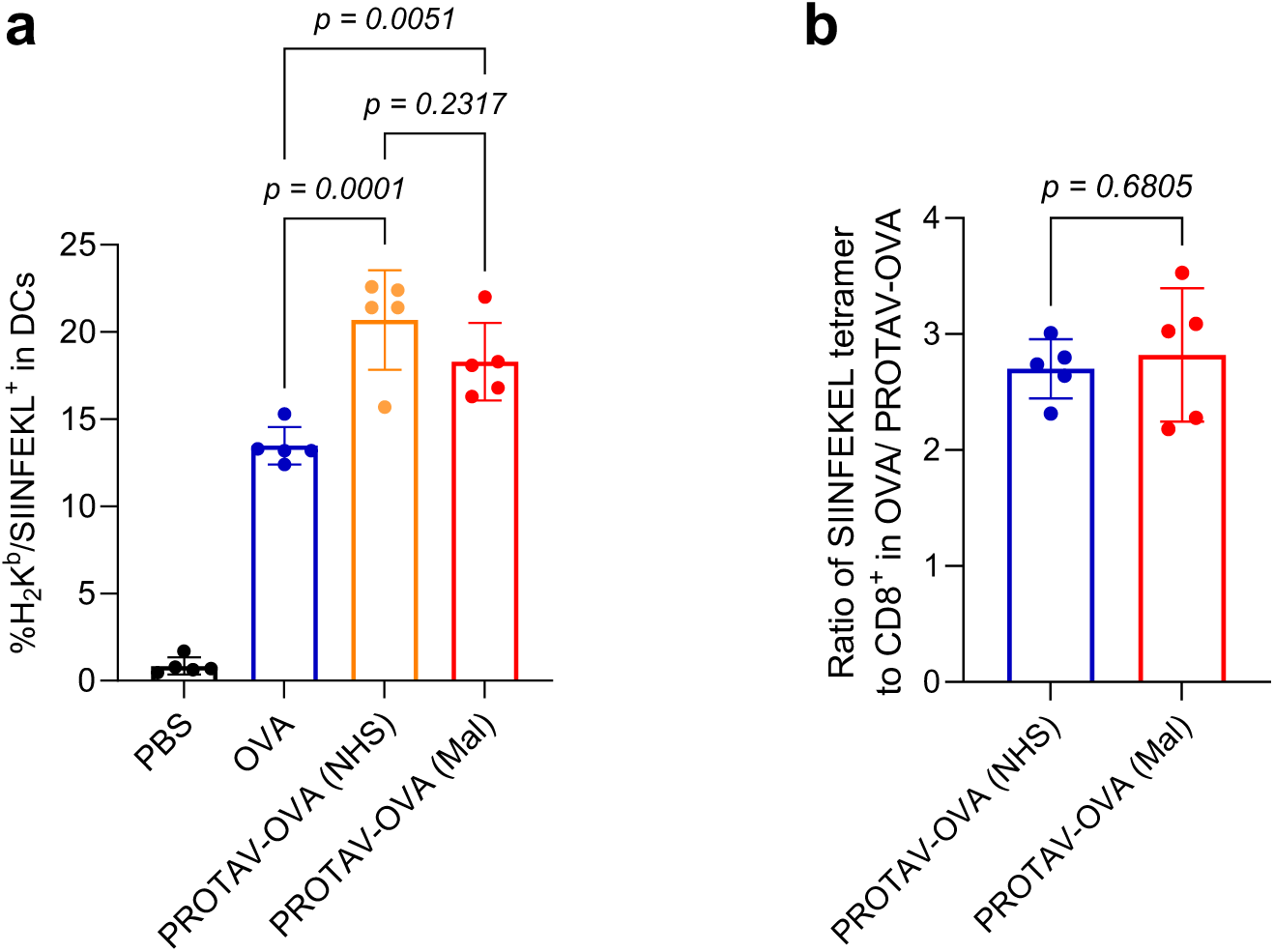
Tetramer staining of mouse PBMC CD8^+^ T cells from mice immunized with PROVA- OVA synthesized using two conjugation methods. **a**, MFI of H-2K^b^/SIINFEKL on DCs quantified from flow cytometry data showing that PROTAV-OVA promoted the presentation of SIINFEKL antigen epitope on DC2.4 cells following a 24-h treatment. **b**, The PROTAV-OVA: OVA ratio of H-2K^b^/SIINFEKL tetramer-positive PBMC CD8^+^ cell fractions elicited by two PROTAV-OVA synthesized by two different conjugation methods (NHS-EDC and maleimide-thiol). These results suggest that these two conjugation chemistries enabled PROTAV-OVA for comparable enhancement of T cell responses over OVA. Mice were immunized by *s.c.* administration at tail base (day 0, day 14). PBMCs were collected for T cell analysis on day 21. CpG (2 nmole) adjuvant was mixed with OVA or PROTAV-OVA (10 μg).

**Supplementary Fig. 7.**
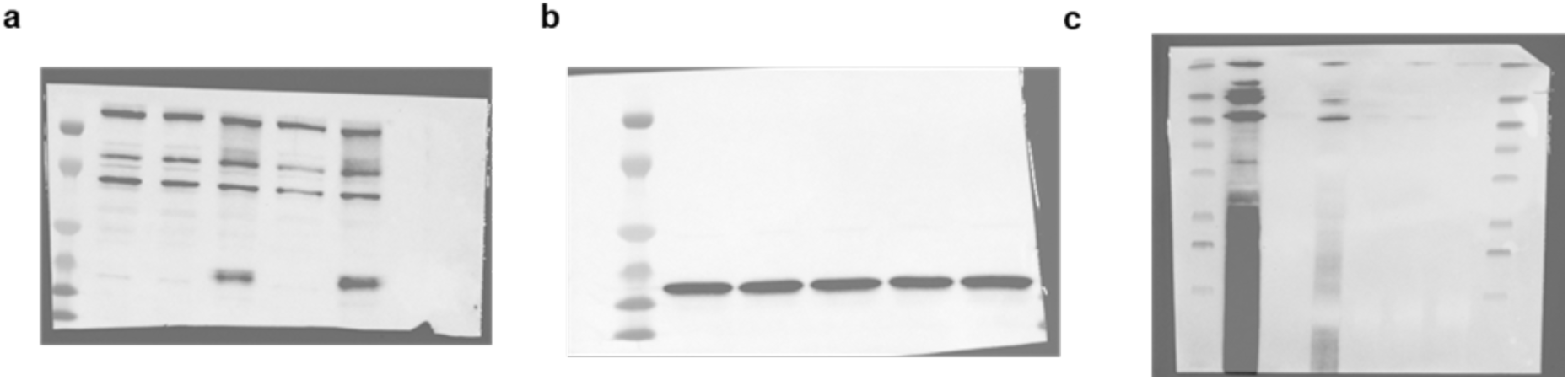
Complete images of Western blot results shown in Fig. 3. (**a, b**) correspond to Fig. 3e, and (c) corresponds to Fig. 3f.

**Supplementary Fig. 8.**
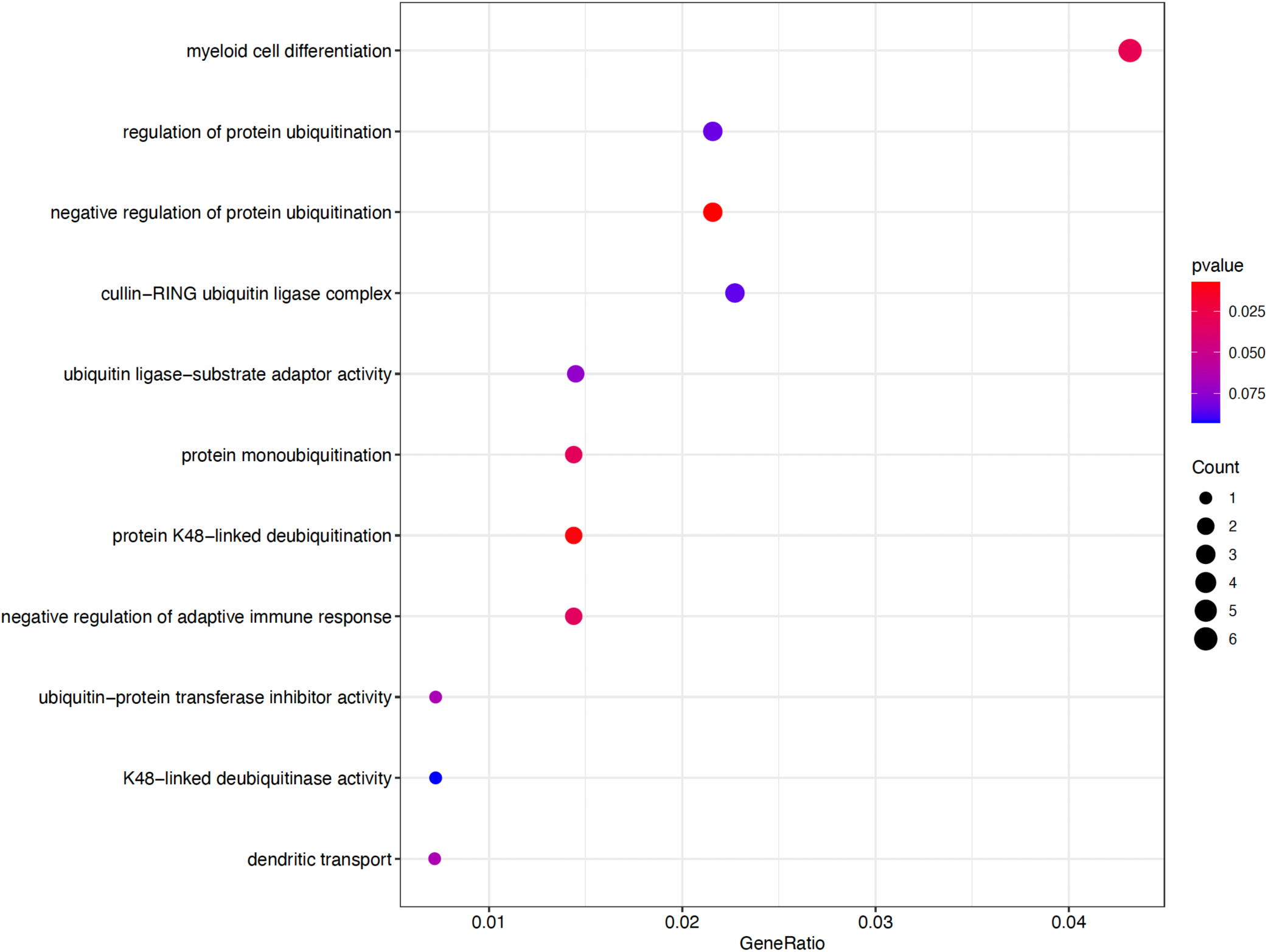
RNA-seq GO analysis of the top 140 differentially expressed genes from BMDCs treated with PROTAV-OVA *vs*. OVA. The results indicate significant enrichment in pathways associated with ubiquitination and immunomodulation in BMDCs treated with PROTAV-OVA relative to OVA. 2 nmole/well CpG was used as an adjuvant, and all RNAseq GO analysis was conducted by using data from the treatment of CpG alone as background. Treatment: 24 h.

**Supplementary Fig. 9.**
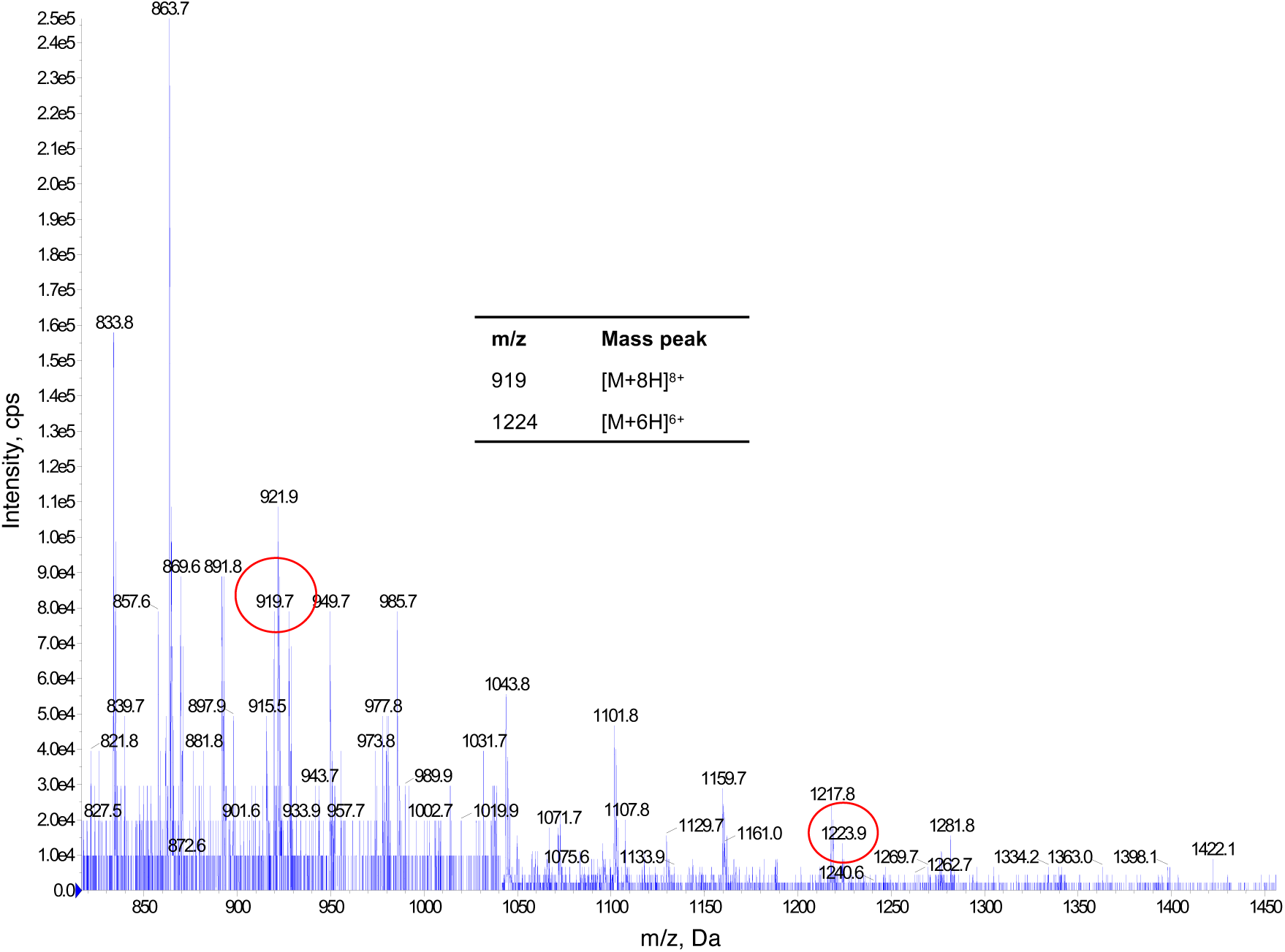
ESI-MS spectra of PROTAV-TgT.

**Supplementary Fig. 10.**
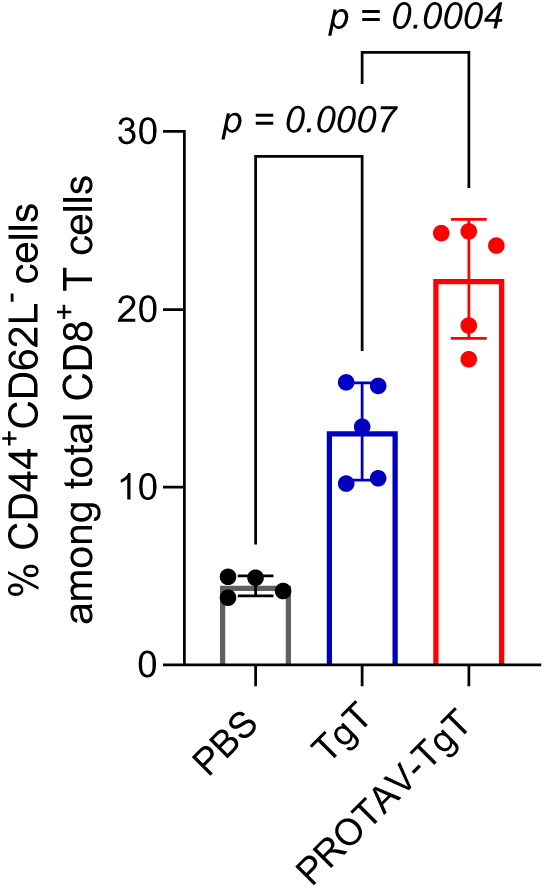
Flow cytometry analysis of CD44 and CD62L levels on PBMC CD8^+^ T cells demonstrates that PROTAV-TgT elicited CD8^+^ T cell memory. Vaccines were delivered by SM-102 LNPs (dose: 20 μg antigen, 2 nmole CpG, 1 nmole Svg3).

**Supplementary Fig. 11.**
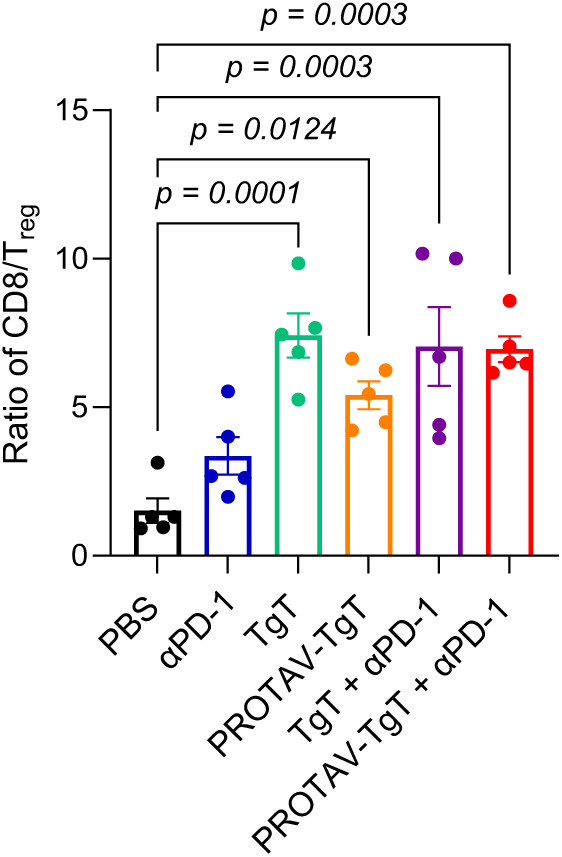
CD8/Treg ratio in B16F10 TME following treatment with PROTAV-TgT and ICB. Vaccines were loaded in SM-102 LNPs (dose: 50 μg antigen, 2 nmole CpG, 1 nmole Svg3) and were *s.c.* injected at mouse tail base. αPD-1: 150 μg, *i.p*. administration.

**Supplementary Fig. 12.**
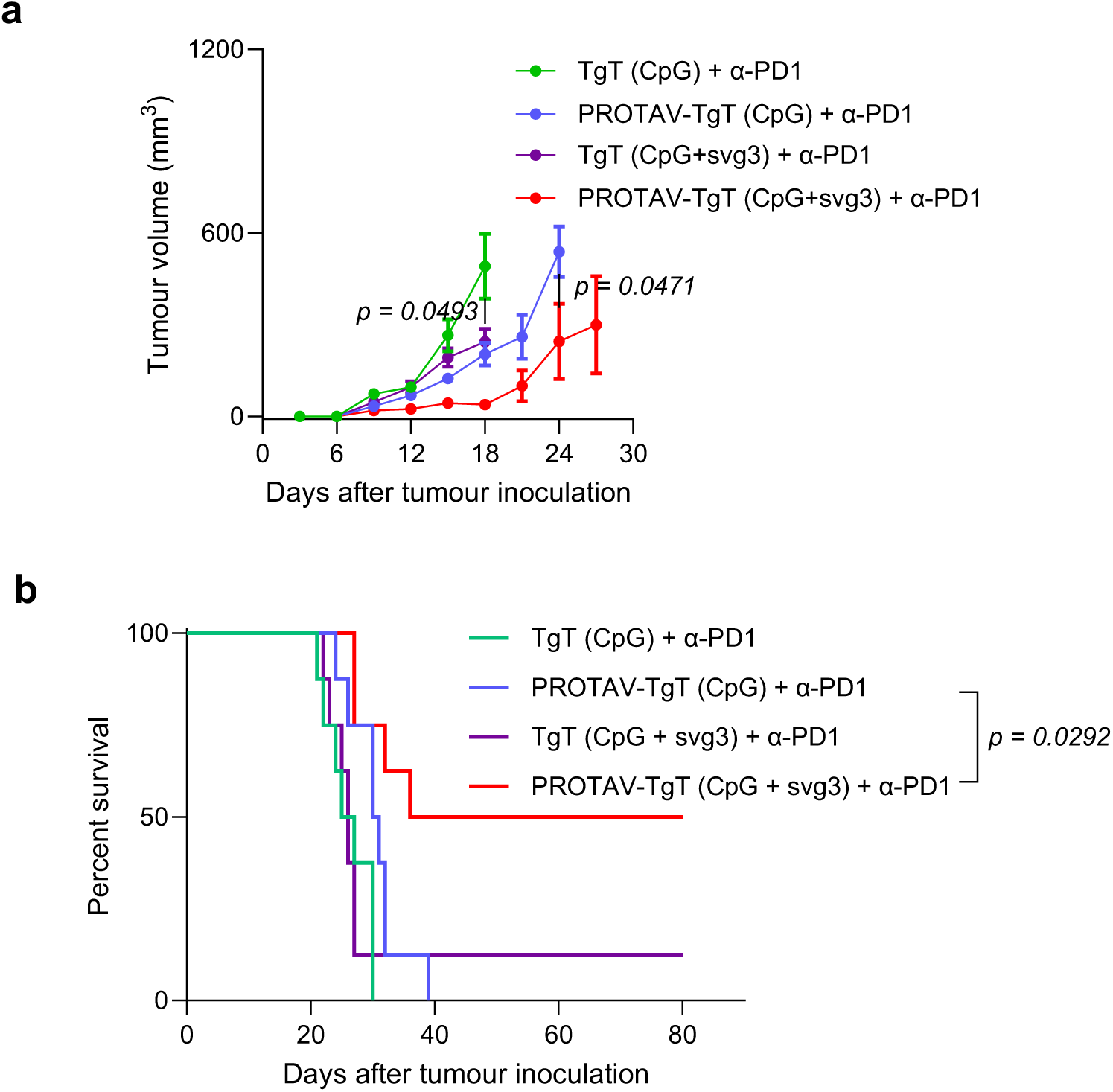
Tumor growth curves (a) and Kaplan-Meier mouse survival curves (b) of B16F10 melanoma-bearing C57BL/6 mice after the indicated αPD-1 combination therapies with PROTAV-TgT or TgT with CpG single adjuvant or Svg3/CpG biadjuvant. PROTAV-TgT with biadjuvant Svg3/CpG outperformed PROTAV-TgT with single adjuvant CpG to inhibit tumor growth. Vaccines were loaded in SM-102 LNPs (dose: 50 μg antigen, 2 nmole CpG, 1 nmole Svg3) and were *s.c.* injected at mouse tail base. αPD-1: 150 μg, *i.p*. administration.

**Supplementary Fig. 13.**
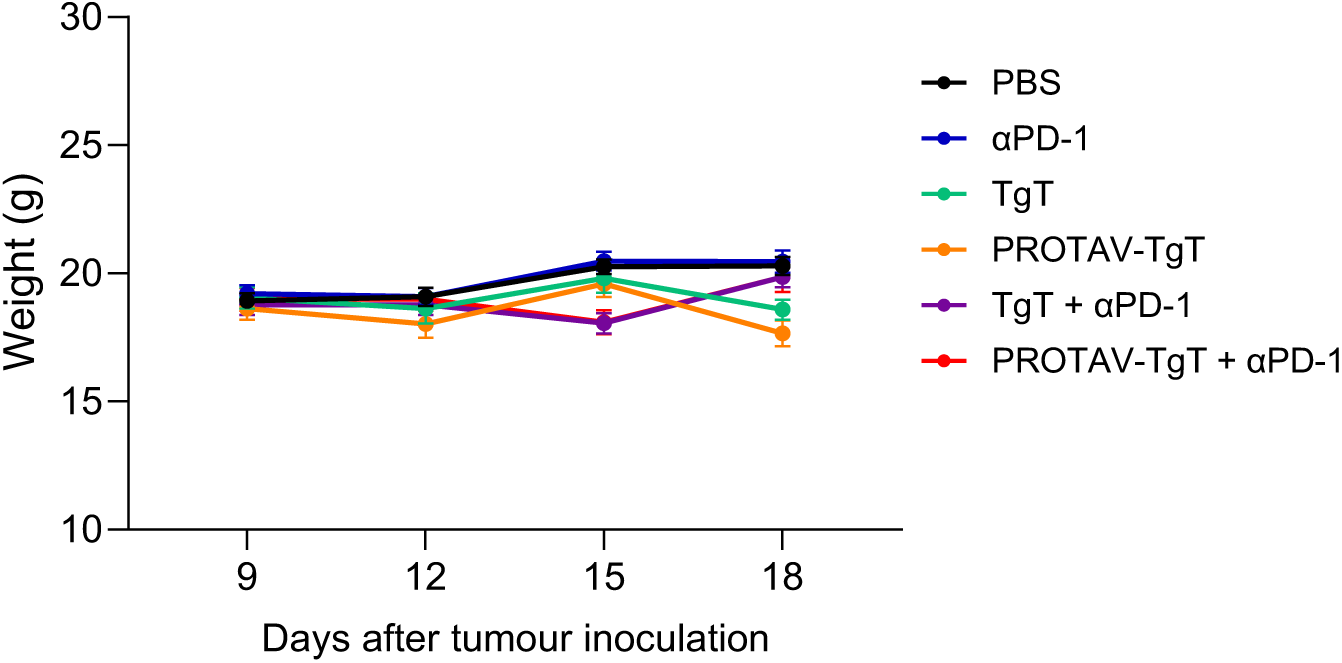
Body weight of B16F10 tumor-bearing mice after treatment with PROTAV + ICB or controls. Vaccines were loaded in SM-102 LNPs (dose: 50 μg antigen, 2 nmole CpG, 1 nmole Svg3) and were *s.c*. injected at mouse tail base. αPD-1: 150 μg, *i.p*. administration.

**Supplementary Fig. 14.**
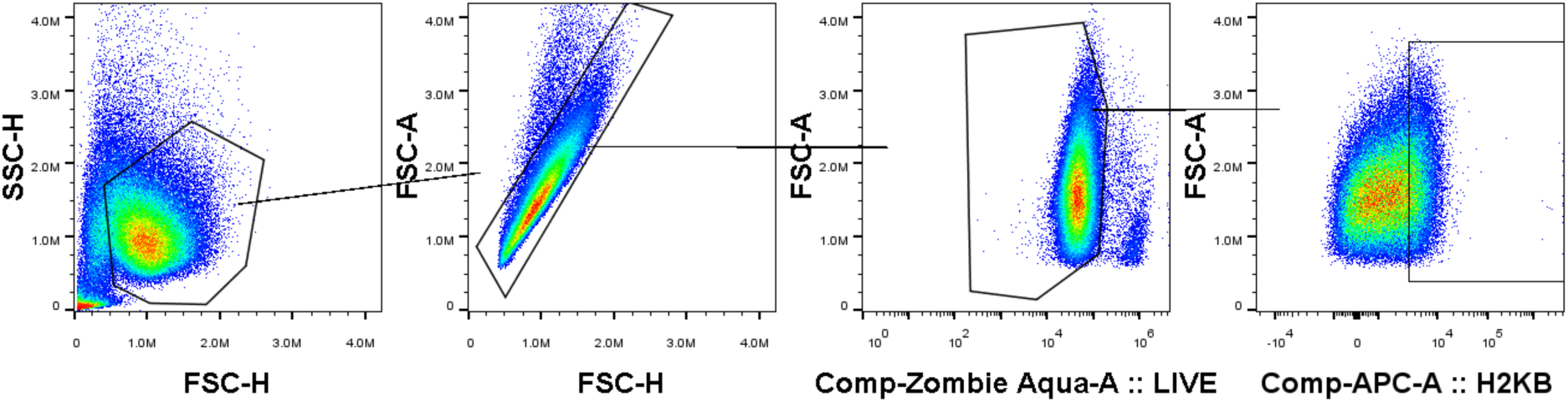
Gating strategy for H-2K^b^/SIINFEKL staining for cultured DCs.

**Supplementary Fig. 15.**
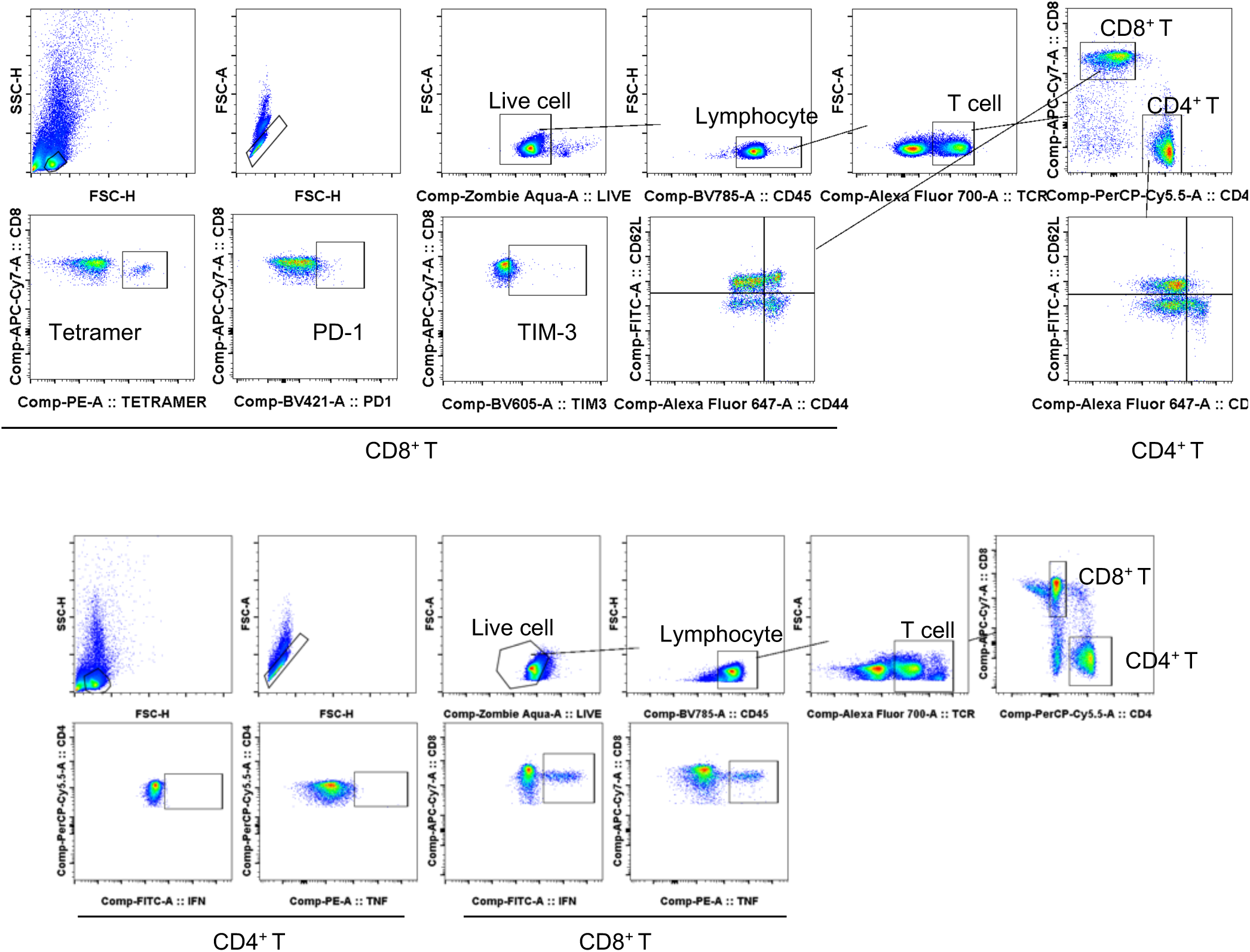
Gating strategy for PBMC T cell staining.

**Supplementary Fig. 16.**
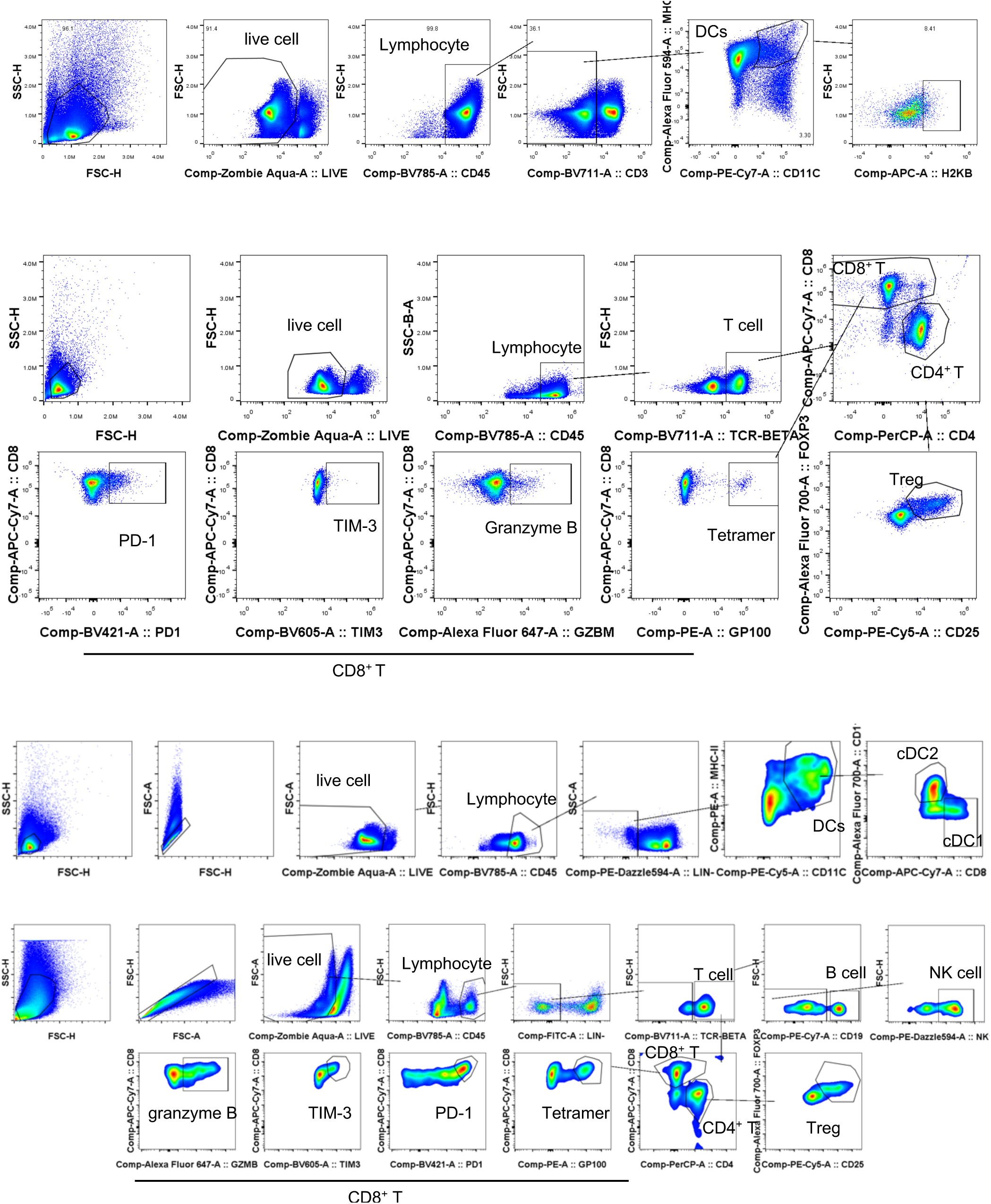
Gating strategy for staining DCs and T cells from lymph nodes.

**Supplementary Fig. 17.**
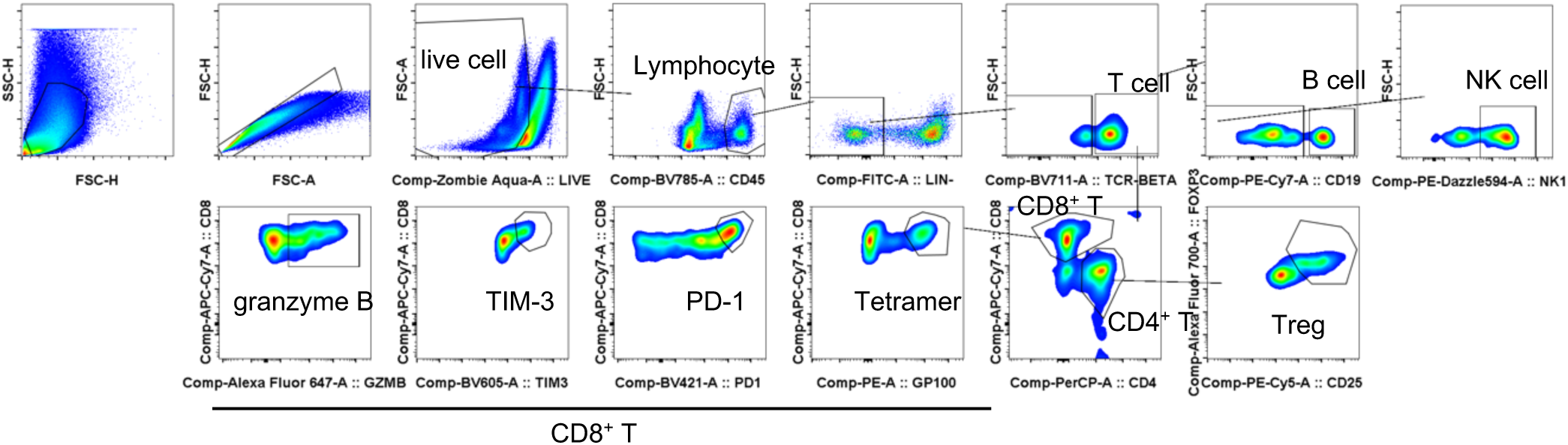
Gating strategy for TME T cell staining.

## Supplementary Tables

**Supplementary Table 1.**
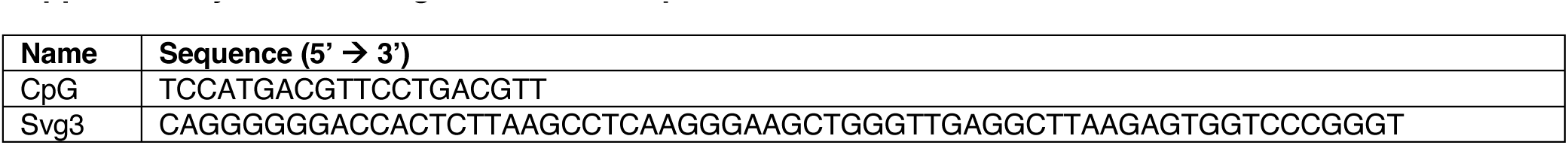
Oligonucleotide sequences.

**Supplementary Table 2.**
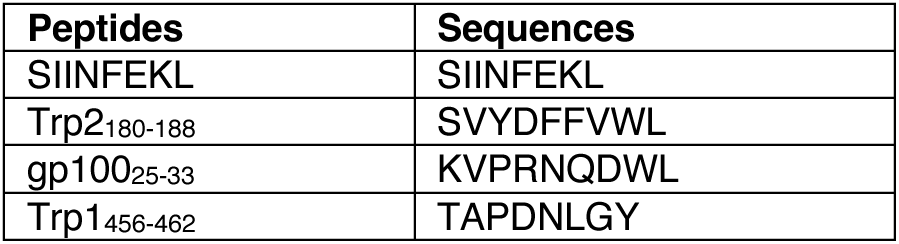
A list of peptides used in this study.

**Supplementary Table 3.** A list of antibodies used in this study.

| Targets | Fluorochromes | Clones | Vendors | Catalogue # |
| --- | --- | --- | --- | --- |
| CD45 | Brilliant Violet 421 | 30-F11 | BioLegend | 103133 |
| CD11c | Alexa Fluor 594 | N418 | BioLegend | 117346 |
| CD11b | FITC | M1/70 | BioLegend | 101205 |
| CD8α | APC-Cy7 | 53-6.7 | BioLegend | 100713 |
| CD4 | PerCP-Cy5.5 | GK1.5 | BioLegend | 100433 |
| CD103 | Brilliant Violet 605 | 2E7 | BioLegend | 121433 |
| CD205 | PE-Cy7 | NLDC-145 | BioLegend | 138209 |
| F4/80 | APC-Cy7 | BM8 | BioLegend | 123117 |
| CD11b | PE-Cy5 | M1/70 | BioLegend | 101209 |
| NK1.1 | APC | S17016D | BioLegend | 156505 |
| CD3 | PerCP-Cy5.5 | 17A2 | BioLegend | 100217 |
| CD279 (PD-1) | Brilliant Violet 421 | 29F.1A12 | BioLegend | 135217 |
| IFN-γ | PE | XMG1.2 | BioLegend | 505807 |
| TNF-α | FITC | MP6-XT22 | BioLegend | 506303 |
| CD44 | Alexa Fluor 647 | IM7 | BioLegend | 103018 |
| CD62L | FITC | MEL-14 | BioLegend | 104405 |
| CD25 | FITC | 3C7 | BioLegend | 101907 |
| FoxP3 | Alexa Fluor 647 | MF-14 | BioLegend | 126407 |
| IL-10 | Brilliant Violet 711 | JES5-16E3 | BioLegend | 505041 |
| MerTK | APC | 2B10C42 | BioLegend | 151508 |
| XCR1 | APC-Cy7 | ZET | BioLegend | 148224 |
| CD19 | PE-Dazzle 594 | 6D5 | BioLegend | 115554 |
| Granzyme B | APC | QA16A02 | BioLegend | 372203 |
| Ubiquitin | N/A | P4D1 | BioLegend | 646302 |
| CD279 (PD-1) | N/A | RMP1-14-CP162 | Bio X Cell | CP162 |
| CD152 (CTLA-4) | N/A | 9H10-CP146 | Bio X Cell | CP146 |
| β-actin | N/A | 15G5A11/E2 | Thermo Fisher | MA1-140 |
| OVA | N/A | N/A | Thermo Fisher | PA1-196 |
| GAPDH | N/A | 5-E10 | Thermo Fisher | MA5-45076 |

